# GLUCURONIDATION: THE LAST DEFENSIVE LINE AGAINST BILE ACID TOXICITY DURING CHOLESTASIS

**DOI:** 10.1101/2025.08.23.671933

**Authors:** Jocelyn Trottier, Audrey-Anne Lavoie, Mélanie Verreault, Ewa Wunsch, Robert J. Straka, Nisanne Ghonem, Piotr Milkiewicz, Olivier Barbier

**Author notes:** Contact information:* Olivier Barbier, Ph. D., FAASLD, Laboratory of molecular pharmacology, CHU de Québec Research Centre, 2705 Laurier Boulevard, Québec (QUE) G1V4G2 CANADA.

## Abstract

Glucuronidation is a detoxification reaction for bile acids (BA). The present study evaluates its efficiency to protect liver cells during acute (biliary obstruction, BO) and chronic (primary biliary cholangitis, PBC and primary sclerosing cholangitis, PSC) cholestatic situations. BA-glucuronides (G) were first profiled in sera from control, PBC, PSC and BO donors. An impressive accumulation of BA-Gs, formed from toxic BAs, is observed in sera from BO patients. Liver samples from control, PBC, or PSC donors and human hepatic cells were exposed to BAs, BA-G, agonists of the BA sensors Pregnane X-Receptor (PXR) or Farnesoid X-Receptor (FXR) and were analyzed for the expression of BA-conjugating UDP-glucuronosyltransferase (UGT) enzymes using quantitative RT-PCR and/or immnunoblot. Expression of the BA-conjugating UGT1A3, 1A4, and 2B4 enzymes was increased in hepatocytes exposed to BA levels mimicking BO. PXR activation also induced expression of the 3 UGTs, while FXR agonists only activated the UGT1A4 and 2B4 genes. Gelshift and luciferase transfection assays identified PXR response elements within UGT1A3 and 1A4 gene promoters. Finally, Inflammation markers have been reduced in hepatocytes treated with BA-G instead of BA.

**Conclusion:** The present study identifies glucuronidation as a self-defense mechanism against BA toxicity during cholestasis. While of limited efficiency by itself, this endogenous mechanism is proposed as a novel pharmacological target in the treatment of chronic cholestatic diseases.

## INTRODUCTION

Cholestasis is defined as an impairment of bile flow (1). Chronic cholestatic diseases include a range of disorders affecting small and large bile ducts and the gallbladder, such as the immune-mediated disorders, e.g., primary biliary cholangitis (PBC) and primary sclerosing cholangitis (PSC) (1, 2). Acute cholestasis includes biliary obstruction (BO) caused by bile duct stones, chronic pancreatitis, or tumors (1). PBC is characterized by chronic lymphocytic inflammation and fibrotic destruction of interlobular bile (2). PSC patients exhibit progressive obliterative fibrosis of intra- and extrahepatic bile ducts (3). In both diseases, fibrosis may lead to cirrhosis, end-stage liver diseases, and the need for liver transplantation (1, 3). While bile flow can be efficiently restored by biliary stenting in BO patients (1), approved PBC therapeutics include the first line therapy ursodeoxycholic acid (UDCA, Ursodiol®), farnesoid X receptor agonists such as obeticholic acid (OCA, Ocaliva®), activators of the peroxisome proliferator-activated receptor (PPAR), and ileal bile acid transporter inhibitors (2). No medical therapy has been approved so far nor has been shown to improve transplant-free survival. Only low URSO doses are used to ameliorate the biochemical parameters of cholestasis (3).

Independent of their cause, the main features of cholestatic and/or autoimmune liver disorders include an accumulation of toxic compounds, especially bile acids (BAs), in the liver and systemic circulation (1, 4–7). BAs play essential roles in cholesterol homeostasis and dietary lipid absorption but are cytotoxic at elevated concentrations (7). In humans, the circulating BAs mainly correspond to the primary cholic (CA) and chenodeoxycholic acids (CDCA) (7, 8). These primary BAs are conjugated with taurine and glycine and conjugated and unconjugated BAs sustain a strong enterohepatic recirculation, through which they can be deconjugated in the intestine and converted into secondary deoxycholic (DCA) and lithocholic acids (LCA) (7, 8). Beyond these deconjugation and dehydroxylation reactions, several bile salt dehydroxylase and hydroxysteroid dehydrogenase enzymes from the microbiota also convert bile acids into amino acid conjugates and oxo-/isomers (9). A significant proportion of BAs is reabsorbed in the intestine and can then be modified back in the liver (10).

These modifications correspond to reconjugation with taurine, glycine, sulfate, and glucuronide. The accumulation of toxic BAs in the cholestatic liver leads to hepatocellular damage, followed by inflammation, apoptotic cell death, and fibrosis (1, 7, 11). BA toxicity is inversely correlated to their hydrophilicity (11–13), and the promotion of less hydrophobic BAs formation has been proposed as a promising therapeutic approach for cholestatic liver diseases (1, 14). On the other hand, BAs also act as signaling molecules and activate nuclear sensors, such as the Farnesoid X-Receptor (FXR) and Pregnane X-Receptor (PXR) which, in hepatocytes, regulate hepatobiliary homeostasis and bile secretion, as well as other excretory functions (15).

Glucuronidation, catalyzed by UDP-glucuronosyltransferase (UGT) enzymes, is a major detoxification pathway for numerous endo-and xenobiotics. This conjugation reaction transfers a highly hydrophilic glucuronide group to hydrophobic substrates, which then become more water-soluble and consequently less toxic (11, 16). Because several investigations revealed that BA-G accumulate in sera, feces, urine or other fluids from patients suffering of various pathologies such as diabetic kidney disease (17), polycystic ovary syndrome (18), intrahepatic cholestasis (19), biliary obstruction (11), PBC and PSC (20) glucuronidation has emerged as potential protective mechanism against bile acid-induced toxicity. Amongst the human functional UGTs, UGT1A3, 1A4, 2B4, and 2B7 have a remarkable capacity to convert BAs into BA-glucuronides (BA-G) *in vitro* (16, 21, 22). Some of these BA-conjugating UGTs were identified as positively regulated PPAR target genes (21, 23–25), and the ability its agonist fenofibrate to increase BA-G levels was further confirmed in PBC and PSC patients (20, 21, 25). The discovery of functional FXR response elements within the promoter regions of some of these genes (26, 27) led to the idea that accumulating BAs may activate their own glucuronidation during cholestasis, thus contributing to a feed-forward reduction of BA toxicity (16, 28). However, because glucuronide (G) conjugates are rarely investigated in clinics, whether such a mechanism effectively occurs has never been validated in humans. In such a context, the present study aimed at investigating whether the circulating BA-G profile may be differentially altered during chronic, i.e., PBC and PSC, and acute, e.g., BO cholestasis and at evaluating the potential benefits for BA detoxification arising from these modifications.

## METHODS

***Materials and expanded methodological descriptions*** are provided in the supplementary Methods (*Supplementary Material-SM1*).

### Ethic Statement

This study received IRB approval from clinical study review boards at the CHU de Québec research centre (*#95-05-14*), the Minnesota fields centre (*#0210M33785*), the Toronto University Health Network (*#05-0670-AE*) and Pomeranian Medical University (Bioethics commission, Pomeranian Medical University in Szczecin, Poland: resolution Nu BN-001/43/06. (4, 5, 21, 29), and conforms to the ethical guidelines of the 1975 Declaration of Helsinki. Informed consent was obtained from each volunteer.

### Subjects

The control, PBC, PSC and BO populations for serum profiling were previously reported (4, 5, 21, 29), and are extensively described in the Supplementary Methods. Briefly, controls were selected among the participants in the Genetics of Lipid Lowering Drugs and Diet Network (GOLDN) study, recruited at the Minnesota fields center (Minneapolis, MN) (21, 29–31). BO patients (8♂, 9♀) were recruited at the Pomeranian Medical University (Szczecin, Poland) (5, 11). PBC patients were from the Toronto Western Hospital (8♀, University Health Network, Toronto, Canada) and the Pomeranian Medical School (4♀) (4). None of the BAs analyzed displayed a significant difference between the two PBC groups (*SM2*). PSC volunteers (5♂, 1♀) were also recruited at the Pomeranian Medical University (4). Because PBC and PSC are gender-related pathologies (14, 32) and circulating BA profiles significantly vary in men and women (21, 29), 3 control groups were selected based on sex and age, for comparison with BO (20♂, 20♀), PBC (27♀, 3♂), and PSC (3♀, 27♂) patients (*SM3&6*) (4, 5). Demographics and liver biochemistry from non-cholestatic volunteers and cholestatic patients have been previously reported (4, 5, 21, 29) and are summarized in *SM3.* Donors (6♂, 6♀) for non-cholestatic liver samples were as previously reported (33), and are extensively described in the *Supplementary methods & SM4*. The study group for liver samples consisted of 22 cholestatic patients: 12 with PBC (2♂, 10♀) and 10 with PSC (8♂, 2♀). Detailed demographic data are shown in *SM5*.

### Bile acid glucuronide measurement

Concentrations of 11 BA-G were determined using high-performance liquid chromatography-tandem mass spectrometry (LC–MS/MS), as previously reported (*Supplementary Methods*) (21). The chromatographic system consisted of an Alliance 2690 HPLC instrument (Waters, Milford, MA), and the tandem mass spectrometry system was an API4000 mass spectrometer (Applied Biosystems, Concord, Canada).

Sera were previously analyzed for a series of 17 unconjugated, taurine-, glycine- or sulfate-conjugated BA species using LC-MS/MS quantification (4, 5). These concentrations were used to calculate the relative abundance (percentage) of BA-Gs and the metabolic ratio (MR; ratio glucuronide *versus* its unconjugated precursor) for each glucuronide.

### Cell culture, mRNA, and protein level determination and glucuronidation assays

Cryopreserved human hepatocytes from 5 donors (*SM7*) were obtained from Celsis-InVitro Technologies, and cultured as described in the *supplementary Methods*. Hepatocytes were treated with vehicle (DMSO, 0.1%, v/v), bile acids (*SM8*) (4, 5, 29), FXR (GW4064, 1 or 5 µM) or PXR (rifampicin, 20 µM) agonists. Human hepatoma HepG2 cells (ATCC) were treated with BAs (50 µM) or BA-Gs, as indicated. Cells were subsequently analyzed for transcript, protein and/or glucuronidation activity levels as detailed in *Supplementary Methods* & (*SM9*).

### Electrophoretic mobility shift assay (EMSA), plasmid cloning, site-directed mutagenesis, and transient transfection assays

EMSA using *in vitro* produced PXR and RXR proteins and the radiolabeled probes listed in *SM9*, were performed as described elsewhere (34–36) and extensively detailed in *Supplementary Methods*.

Luciferase reporter plasmids for the wild type (WT) or mutated (MT), UGT1A3 and/or 1A4 PXR response elements (PXREs) were obtained and transiently transfected into HepG2 cells as detailed in Supplementary Methods & *SM9*.

### Data analyses

BA-G levels were calculated as mean±standard deviation (SD). BA-G concentrations did not satisfy the normal distribution according to the Shapiro-Wilk test, thus the Wilcoxon/Mann-Whitney rank-sum test was used for statistical analyses of changes in BA-G profiles (JMP V7.0.1, SAS Institute Inc, Cary, NC). The statistical significance of differences in mRNA levels and BA-G formation was determined through the Student *t-*test using the JMP V7.0.1 program (SAS Institute Inc.).

## RESULTS

### Circulating bile acid glucuronide profiles are differentially affected in primary biliary cholangitis, primary sclerosing cholangitis, and biliary obstruction sera

BA-G profiles and changes are illustrated in **Figure 1**, and detailed in *SM10-12*. When compared to their respective controls, total BA-G levels were significantly increased in PBC (**Figure 1A**, *p*<0.05) and BO patients (**Figure 1C**, *p*<0.001), but not in PSC samples (**Figure 1B**). However, the relative abundance of glucuronide conjugates within the total circulating BA pool sustained opposite changes with a significant (*p*<0.001) reduction in BO samples when compared to controls (**Figure 2A**, *SM12*). This may reflect the huge accumulation of non-glucuronidated BAs (>58 times) previously reported in these sera (5). A similar trend was also observed in PSC samples, but not in PBC sera (**Figure 2A**, *SM10-11*).

**Figure 1.**
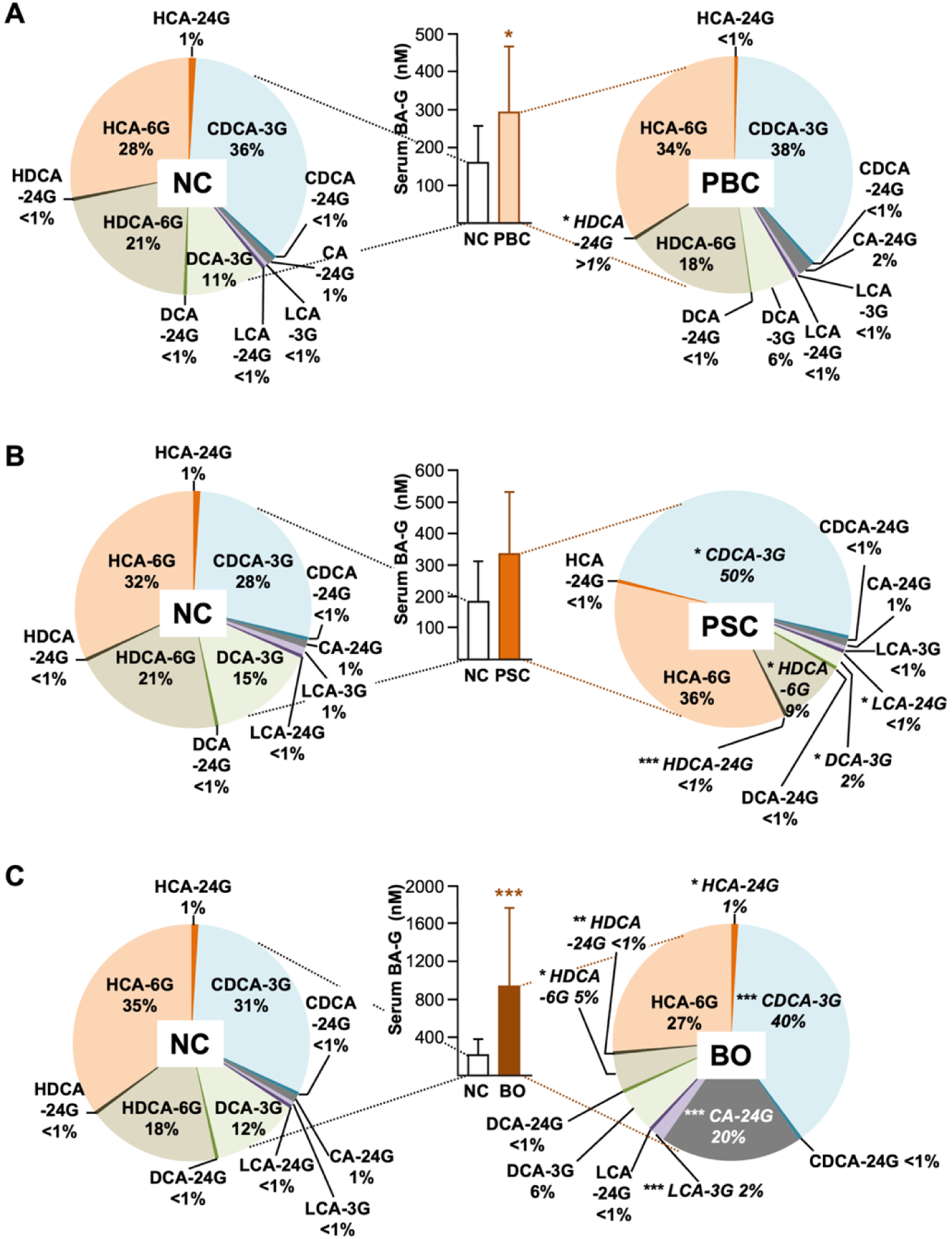
Biliary obstruction (BO), but not primary biliary cholangitis (PBC) or primary sclerosing cholangitis (PSC), causes major changes in the circulating bile acid-glucuronide profile. Serum samples from non-cholestatic volunteers (NC), PBC (**A**), PSC (**B**), and BO (**C**) patients were analyzed for their contents in 11 bile acid-glucuronide (BA-G) species using liquid chromatography-tandem mass spectrometry. (**A-C**) Histograms represent the mean±SD of the serum BA-G levels as determined as the sum of all glucuronide species. Pie charts represent the relative proportion (expressed as percentage) of each BA-G species within the total glucuronide pool, i.e., sum of all BA-G. Because PBC and PSC are gender-related pathologies, and because circulating profiles of BA-Gs significantly differ in men and women, 3 control groups were selected based on sex and age, for comparison with BO, PBC, and PSC patients. *P*-values were determined by the Wilcoxon/Mann-Whitney rank-sum test: *p<0.05; **p<0.01; ***p<0.001. The complete results are provided in supplemental materials 10 to 12.

**Figure 2.**
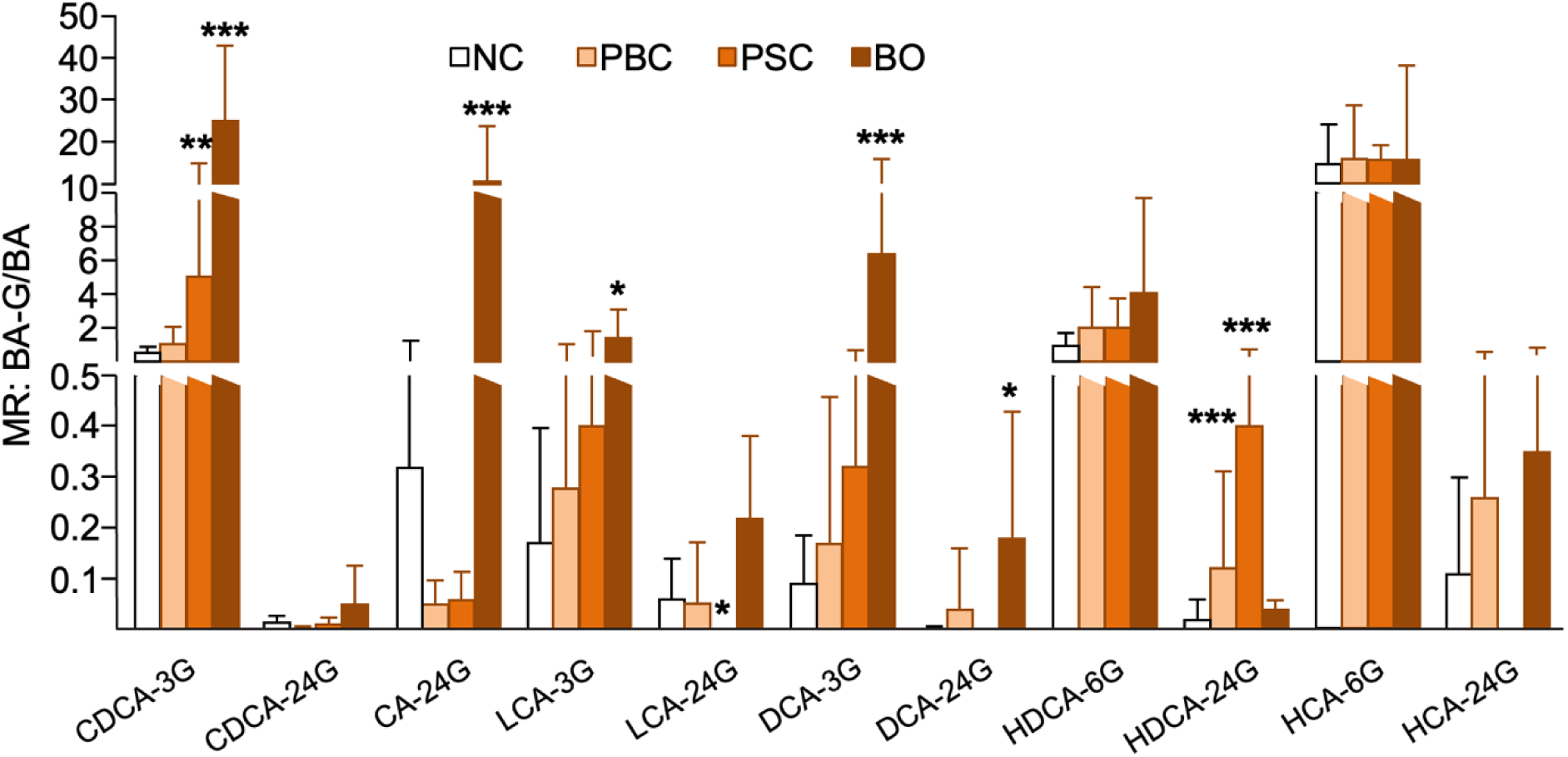
Toxic bile acid species are preferentially glucuronidated during biliary obstruction. Serum bile acid glucuronides were determined through LC-MS/MS measurements in samples from non-cholestatic volunteers (NC), PBC (**A**), PSC (**B**), and BO (**C**) patients (pt). The metabolic ratio for each species was calculated as the ratio of glucuronide *versus* its unconjugated precursor. Statistically significant differences were determined using the Wilcoxon/Mann-Whitney rank-sum test: *p<0.05; **p<0.01; ***p<0.001. The complete results are provided in supplemental materials 10 to 12.

Analyses of the BA-G pool composition revealed major differences between PBC, PSC, and BO samples (**Figure 1**). In PBC, the unique significant change corresponded to the 3.9-fold accumulation of HDCA-24G (p<0.001) (**Figure 1A**, *SM10*). PSC patients presented significant increases in CDCA-3G (*p*<0.05) and HDCA-24G (*p*<0.001) and decreases in LCA-24G (*p*<0.05), DCA-3G (*p*<0.05) and HDCA-6G (*p*<0.05) levels, (**Figure 1B**, *SM11*). Consequently, the proportion of CDCA-3G within the BA-G pool increased from 28% (controls) to 50% (PSC), while those of HDCA-6G and DCA-3G were 2.9- and 8.2-times reduced (p<0.05), respectively (**Figure 1B**). The most impressive changes were observed in samples from patients with biliary obstruction, where the 87.8-, 10.1- and 5.5-fold increases in CA-24G (*p*<0.001), LCA-3G (*p*<0.001) and CDCA-3G levels (*p*<0.001), respectively (*SM12*), resulted in drastic modifications of the BA-G pool. Indeed, CA-24G, a minor component in control samples (1%) was the 3^rd^ most abundant (20%) BA-G in BO sera, while the representation of DCA-3G and HDCA-6G decreased from 12 and 18% (controls) to 6 and 5% (BO), respectively (**Figure 1C**). Consequently, the contribution of derivatives formed from primary acids (i.e. CA- and CDCA-glucuronides) increased from 32.2±17.7% in controls to 72.6±23.3% in BO. By contrast, the proportion of non-toxic 6α−hydroxylated BA-G (HCA- and HDCA-G) which accounted for 51.1±20.3% of glucuronides in non-cholestatic samples was reduced to 17.6±22.1%.

### Cholestasis changes the selectivity of glucuronidation for BA substrates

To further investigate whether changes in BA-G pool composition reflect modifications of the glucuronidation capabilities or an increased availability for glucuronide conjugation, we next calculated the metabolic ratio for each conjugate over their respective parent BA (**Figure 2**). When compared to their respective controls, only PBC samples exhibited an increased HDCA-24G/HDCA ratio (*p*<0.001), while PSC sera displayed higher MRs for CDCA-3G (*p*<0.01) and HDCA-24G (*p*<0.001), and a lower LCA-24G ratio (*p*<0.05) (**Figure 2**). As above, the most spectacular changes were observed in samples from BO patients where DCA-3G (*p*<0.001), CA-24G (*p*<0.001), DCA-24G (*p*<0.05), CDCA-3G (*p*<0.001) and LCA-3G (*p*<0.05) ratios were respectively 78.0-, 58.7-, 55.8-32.4- and 13.1-fold increased (**Figure 2**). By contrast, none of the MR values for the four 6α-hydroxylated BAs investigated were significantly affected by BO (**Figure 2**).

### BAs stimulate the hepatic expression of UGTs involved in their own glucuronidation

To evaluate the possibility that BAs self-modify their hepatic glucuronidation, we exposed human hepatocytes to the level of BAs found in sera from non-cholestatic (4, 5), PBC (4), PSC (4) or BO (5) donors (*SM8*). BA levels from PBC caused significant accumulations of UGT1A3, 1A4, and 2B4 transcripts, while BA concentrations mimicking PSC also favored UGT1A3 and 1A4 mRNA expression (**Figure 3A**). However, the highest mRNA inductions were observed for UGT1A3 (4.8-fold, *p*<0.001), 1A4 (4.5-fold, *p*<0.001), and 2B4 (4.3-fold, *p*<0.001) in cells exposed to the circulating bile acid profile from patients with biliary obstruction (**Figure 3A**). Interestingly, none of the BA conditions affected UGT2B7 mRNAs (**Figure 3A**). Western-blot experiments revealed that conditions mimicking BO also caused accumulation of UGT1A and UGT2B proteins in hepatocytes (**Figure 3B**), while cells exposed to the BA profiles corresponding to PBC and PSC samples exhibited less consistent changes in their UGT protein contents (**Figure 3B**). Accordingly, western-blot analyses of non-cholestatic control (n=12), PBC (n=12), and PSC (n=10) liver extracts suggested that UGT1A and 2B proteins do not accumulate under such chronic cholestatic conditions (**Figure 3C**). Indeed, the UGT2B protein content tended to decrease in liver samples from PSC donors (**Figure 3C**). Unfortunately, the lack of similar samples from patients suffering from BO impaired comparable analyses.

**Figure 3.**
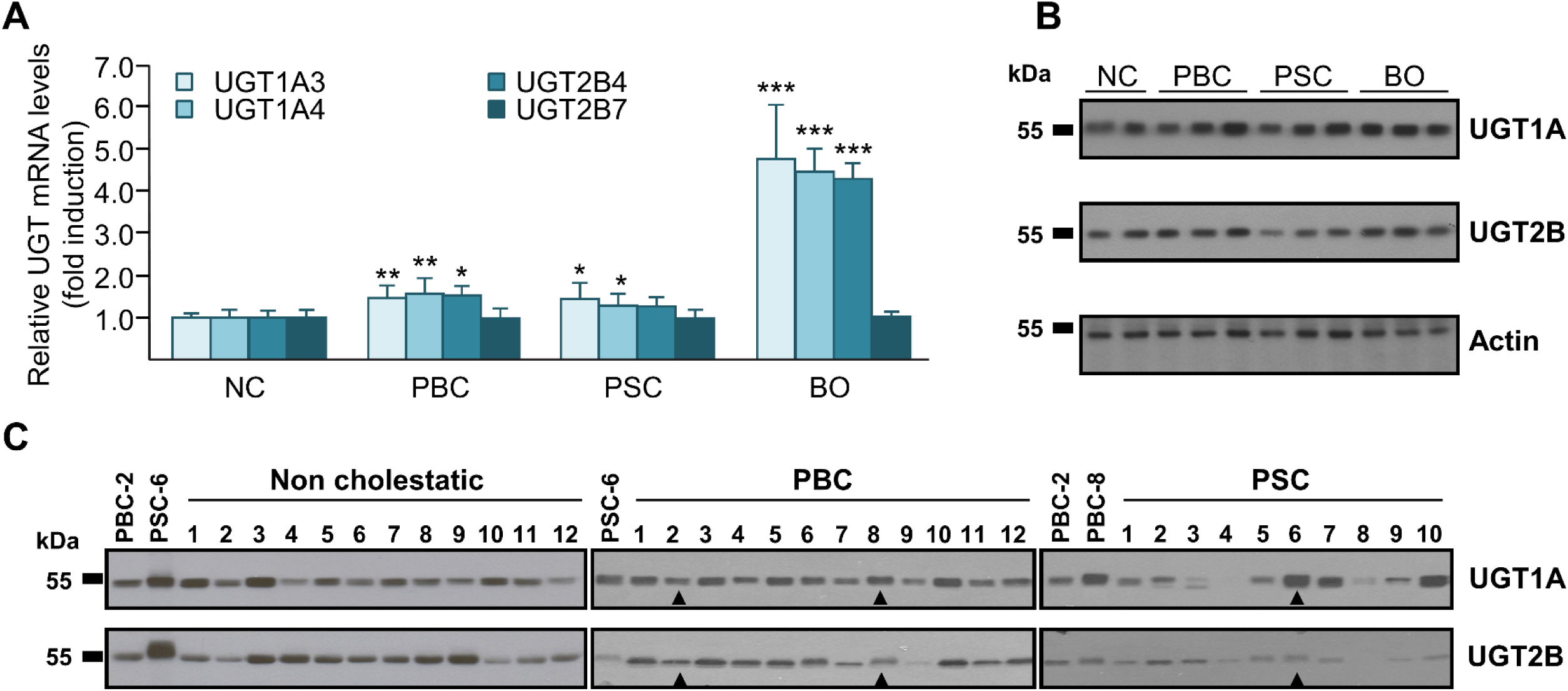
Bile acids differentially affect the expression of the human bile acid-conjugating UGTs in human hepatocytes (A & B), while UGT1A and UGT2B protein levels are barely affected in liver samples from patients with primary biliary cholangitis (PBC) and primary sclerosing cholangitis (PSC) (C). (**A&B**) Primary human hepatocytes were cultured for 24H in the presence of bile acid mixtures mimicking the circulating BA profiles previously detected in non-cholestatic volunteers (NC, controls), PBC, PSC, or biliary obstructed (BO) patients. (**A**) UGT2B4, 2B7, 1A3, and 1A4 mRNA levels were quantified from total RNA through quantitative RT-PCR analyses and normalized with the housekeeping RNA 28S. Values (mean±SD) are expressed relatively to control (NC) set at 1. Statistically significant differences between control and treated cells are indicated by asterisks (Student *t* test: *: p<0.05; **: p<0.01; ***: p<0.001). (**B&C**) The UGT1A and UGT2B protein contents in BA-treated hepatocytes (10 µg) or liver microsomes (10 µg) from non-cholestatic, PBC or PSC donors were evaluated after size separation on SDS-PAGE and hybridization with the anti-UGT1A (1:2,000) and anti-UGT2B (1:2,000) antibodies as indicated. Membranes with samples from BA-treated hepatocytes (**B**) were subsequently hybridized with an anti-actin antibody to ensure the equal loading of each lane. Arrowhead on panel **C** identifies samples used as markers for immunoblot exposure of multiple membranes.

Next, the contribution of the BA sensors in the BA-induced activation of UGT expression and/or activity, was evaluated using primary isolated human hepatocytes cultured in the presence of high-affinity FXR (GW4064) (37) and PXR (rifampicin) (38) agonists (**Figure 4A**). While UGT1A3 and 2B7 mRNA levels remained unchanged, treatment with GW4064 (1 and 5 µM) caused a significant, but limited increase in UGT1A4 transcripts (**Figure 4A**). According to previous reports (24), this FXR agonist caused a dose-dependent induction of UGT2B4 mRNA expression (**Figure 4A**). Such induction also increased UGT2B protein levels (**Figure 4B**). Exposure to rifampicin (20 µM) resulted in UGT1A3 and 1A4 mRNAs accumulation in human hepatocytes from the 2 different donors (**Figure 4C**), which led to a convincing UGT1A protein accumulation (**Figure 4D**). Interestingly, the same treatment increased UGT2B4 transcripts by 21.7-fold, but only in cells from donor 2, and a 47% reduction of UGT2B7 mRNA expression was found in hepatocytes from donor 3 (**Figure 4C**). These changes in UGT expression resulted in an improved formation of BA-G by hepatocytes from donors 4 and 5 (**Figure 4E**). Of the 11 glucuronides investigated, the production of LCA-3 and -24G and HDCA-6 and 24G was significantly increased in treated cells from both donors, while HCA-6G formation remained not significantly changed in all of the cells tested (**Figure 4E**, *SM13*). Other BA-G were more abundantly formed in cells from at least 1 donor.

**Figure 4.**
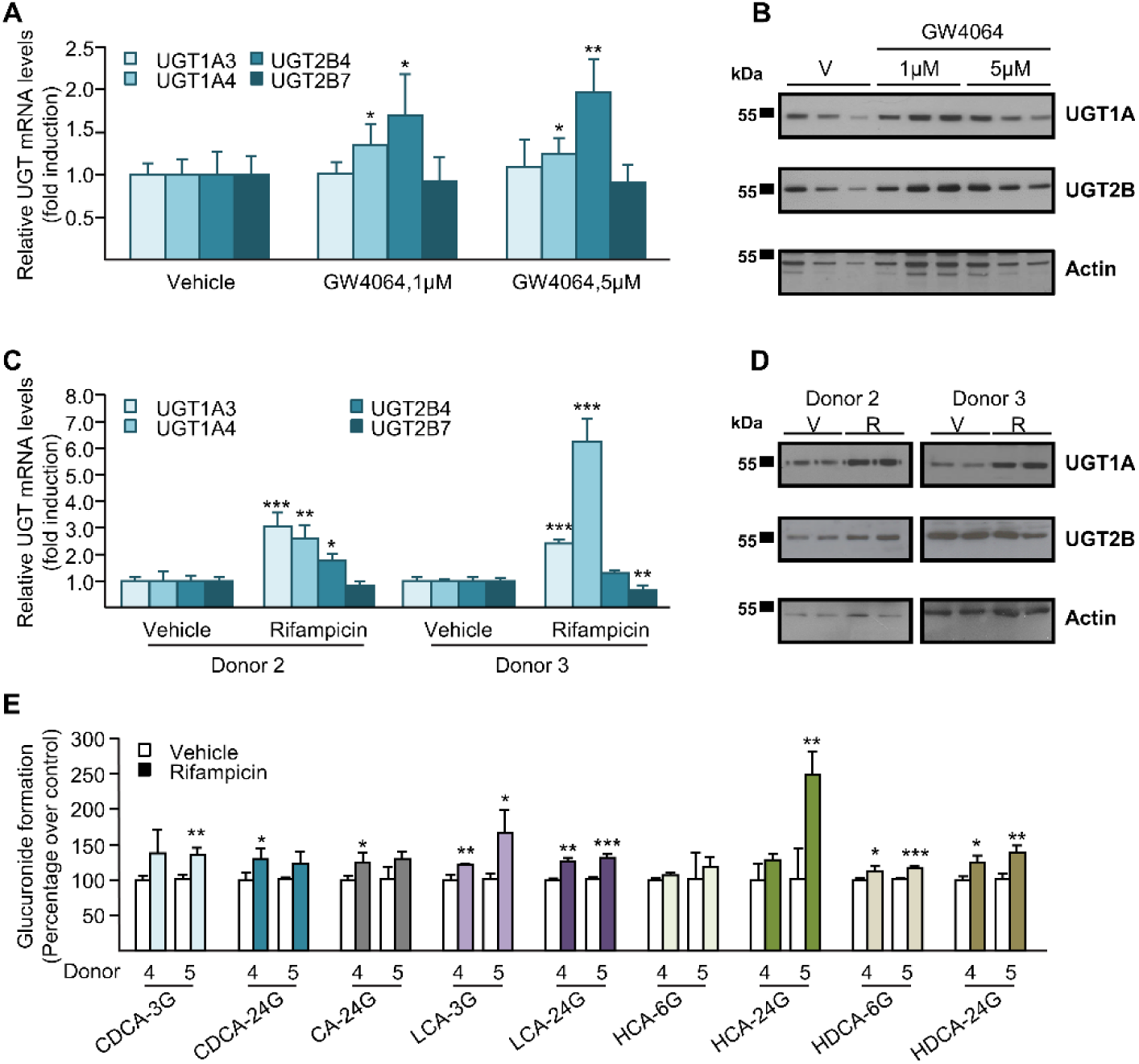
The FXR (GW4064, A&B) and PXR (Rifampicin, C&D) agonists differentially affect the expression and/or activity of human bile acid-conjugating UGTs in hepatocytes. Primary human hepatocytes were treated for 24 (**A-D**) or 48H (**E**) with DMSO (vehicle), 1 and 5 µM GW4064 (donor 1) or 20 µM Rifampicin (donors 2-5). (**A&C**) UGT2B4, 2B7, 1A3, and 1A4 mRNA levels were quantified from total RNA through quantitative RT-PCR analyses and normalized with the housekeeping RNA 28S. Values (mean±SD) are expressed relative to vehicle set at 1.0. Statistically significant differences between control and treated cells are indicated by asterisks (Student *t*-test: *: p<0.05; **: p<0.01; ***: p<0.001). (**B&D**) UGT1A and UGT2B protein levels were visualized in cell proteins (10 µg) through immunoblotting using the anti-UGT1A (1:2,000 dilution) and anti-UGT2B (1:2,000 dilution) antibodies. The same membranes were subsequently hybridized with an anti-actin antibody (1:2,000 dilution) to ensure the equal loading of each lane. (**E**) Bile acid glucuronidation was analyzed through *in vitro* assays performed for 1H in the presence of 100µM BAs, 1 mM UDPGA and 5-10µg HH homogenates. The formation of BA-G was resolved through LC-MS/MS. Values (mean±SD) are expressed relatively (percentage) to control cells. Statistically significant differences between control and treated cells are indicated by asterisks (Student *t*-test: *: p<0.05; **: p<0.01; ***: p<0.001). The complete results are provided in supplemental materials 13.

### PXR binds to and activates response elements in the UGT1A3, 1A4 and 2B4 promoter genes

FXR response elements within UGT gene promoters were previously identified (26, 27), and we next investigated the mechanisms of PXR to modulate UGT1A3, 1A4, and 2B4 mRNA expression. Computer-assisted analysis (39) of the UGT1A3 and 1A4 promoter regulatory regions revealed the presence of various PXR binding motifs (40) corresponding to degenerated direct (DR) or everted repeat (ER) of the AGGTCA hexamer sequences separated by 3 (DR3), 4 (DR4) or 6 (ER6) nucleotides (*SM14*). EMSAs revealed that only the PXREs located at positions -8,056bp, -6,930bp, -1,935bp, and -260bp of the UGT1A3 promoter upstream of the transcription start site were complexed with the PXR/RXR heterodimer (*SM15*). Similar experiments performed with the UGT1A4 promoter revealed that only the -2,014bp DR3-like and -260bp DR4-like sequences were bound by this heterodimer (*SM15*). Interestingly, these 2 PXREs are 100% identical to those found at positions -1,935bp and - 260bp of the UGT1A3 promoter (*SM14*). Additional investigations (**Figure 5**), show that assays with artificially mutated PXREs (line 14), competition analyses with the PXRE from CYP3A4(34) (lines 5-7), unlabeled WT (lines 8-10) or mutated PXREs (lines 11-13), as well as supershift experiments with the anti-PXR antibody (line 4) confirmed the PXR-dependent formation of all these protein-DNA complexes (**Figure 5A-D**). The functional consequences of PXR binding to these response elements were next tested by transfecting HepG2 cells with the TKpGL3 luciferase reporter vector driven by 3 copies of each wild type or mutated PXRE in the presence of PXR and RXR (**Figure 5A-D**). In all cases, activities of UGT1A3 and/or 1A4 PXREs were increased in the presence of rifampicin, an effect lost when the PXR/RXR binding was disrupted through the introduction of mutations within PXREs (**Figure 5**). Finally, a similar experimental strategy allowed the discovery of a functional and positive PXR response element located at position -490bp of the UGT2B4 promoter (**Figure 5E**).

**Figure 5.**
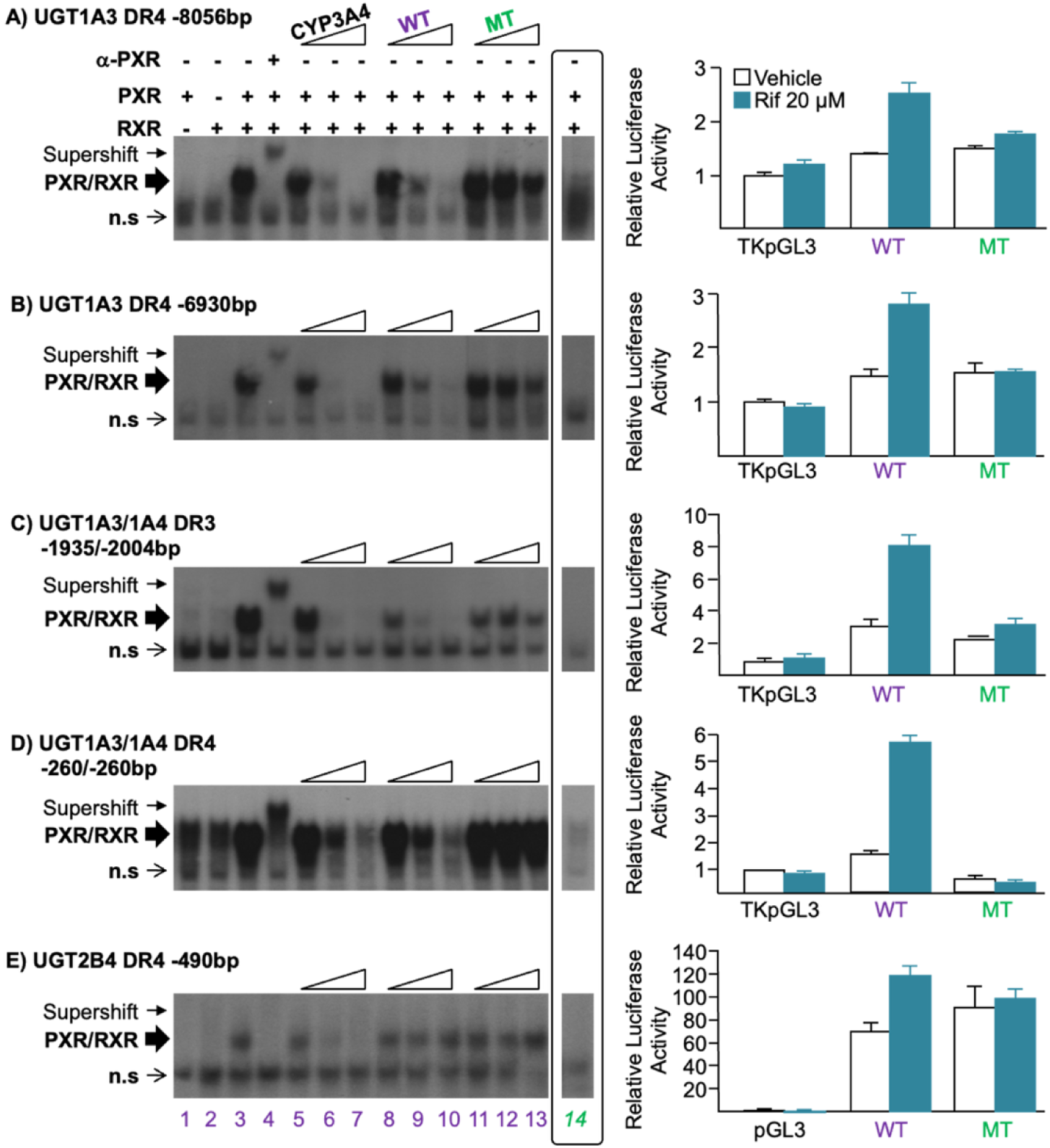
PXR binds to and activates PXREs in the UGT1A3, 1A4, and 2B4 gene promoters. (**A-E, left panels**) Electrophoretic Mobility Shift Assays were performed with end-labeled probes encompassing the wild type (lines 1-13) or mutated (line 14) sequences of the UGT1A3 DR4-8056 bp, UGT1A3 DR4-6930 bp, UGT1A3/1A4 DR3-1935/2004 bp, UGT1A3/1A4 DR4-260 bp or UGT2B4 DR4-490 bp PXREs in the absence or presence of *in vitro* produced PXR, RXR or both PXR and RXR proteins as indicated. Competitions (lines 5-13) on these probes were performed by adding 1-, 10-, or 50-fold molar excess of the indicated cold competitor oligonucleotides in EMSA with RXR and PXR. Supershift experiments (line 4) were carried out using an anti-PXR antibody (α-PXR; 0.2 µg). (**A-D, right paanels**) HepG2 cells were transfected with the indicated plasmids containing 3 copies of the wild type (WT) or mutated (MT) UGT1A3 and/or UGT1A4 response elements upstream of the thymidine kinase (TK) minimal promoter-driven luciferase reporter (TKpGL3), and pRL-NULL in the presence of PXR/RXR. (**E, right panel**) HepG2 cells were transfected with the human UGT2B4p-524 promoter-driven luciferase (Luc) reporter plasmid containing either the wild type or mutated sequences of the -490 bp DR4 element and pRL-NULL. (**A-E, right panels**) Cells were subsequently treated with Rifampicin (RIF, 20 µM) or vehicle (DMSO) for 24H. Values (means±SD) were normalized to internal renilla activity, and expressed as fold-induction of controls (pGL3 or TKpGL3) set at 1.

### Glucuronidation protects against the pro-inflammatory effects of primary and secondary acids

Recent reports indicate that BAs activate hepatic inflammation during obstructive cholestasis (41, 42). To further examine the physiological consequences of the improved BA glucuronidation, HepG2 cells were exposed to unconjugated CDCA, CA, LCA, and HDCA or their glucuronide derivatives (**Figure 6**), and the mRNA expression of the inflammatory markers interleukin (IL)-8, plasminogen activator inhibitor (PAI)-1 and intercellular adhesion molecule (ICAM)-1 were determined (41). ICAM-1 mRNA expression was significantly up-regulated in cells cultured with CDCA, CA, and HDCA (**Figure 6A**). Similarly, PAI-1 mRNAs accumulated in all BA-cultured cells (**Figure 6C**), while only CDCA and LCA caused significant accumulations of IL-8 transcripts (**Figure 6B**). Interestingly, these effects were all significantly reduced when BAs were replaced by their glucuronide conjugates, thus indicating that BA-Gs present lower pro-inflammatory effects than their unconjugated precursors.

**Figure 6.**
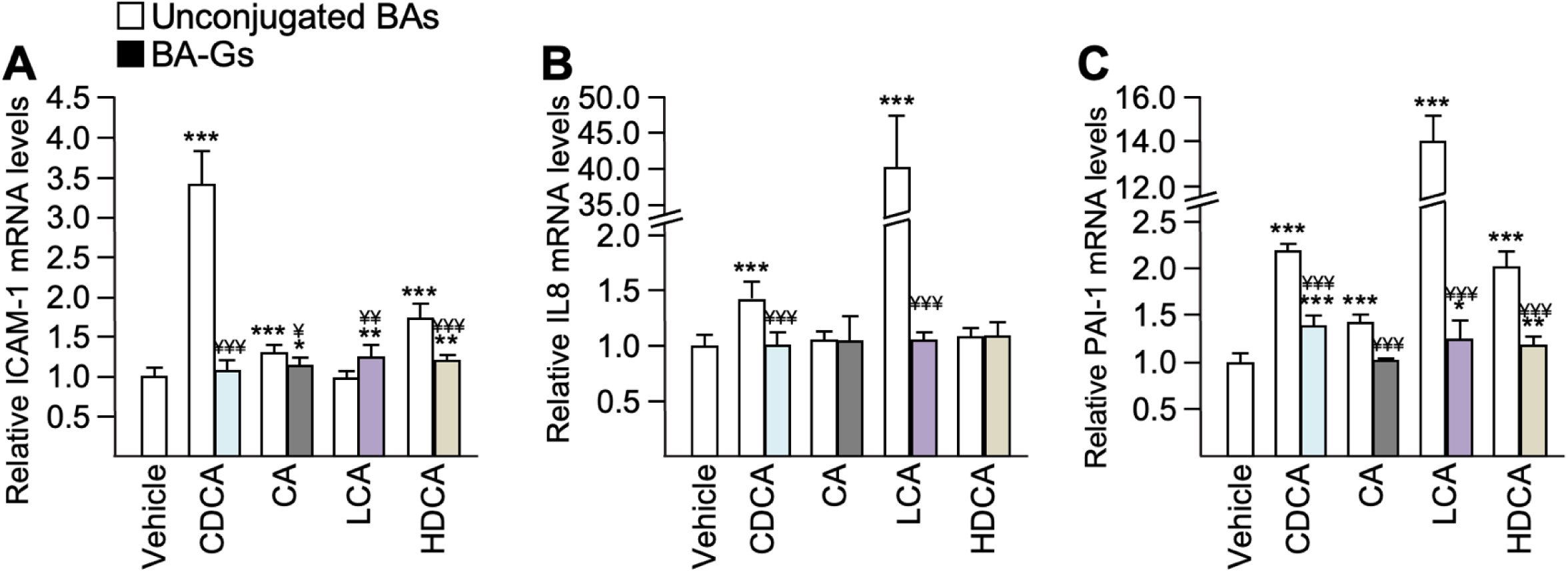
In contrast to their unconjugated precursors, glucuronide conjugates of primary and secondary bile acids fail to activate proinflammatory mediators. Human hepatoma HepG2 cells were cultured for 24H in the presence of DMSO (vehicle) or 50 µM unconjugated CDCA, CA, LCA, and HDCA (white bars) or glucuronidated CDCA-3G, CA-24G, LCA-3G and HDCA-6G (color bars) bile acids. The mRNA levels of ICAM-1(**A**), IL8 (**B**), and PAI-1 (**C**) were measured by quantitative RT-PCR, and normalized with the housekeeping RNA PUM-1 and expressed relative to control (vehicle) set at 1. All values represent the means±SD. Statistically significant differences in vehicle versus treated cells (*p<0.05; **p<0.01; ***p<0.001) and in glucuronide-versus unconjugated-exposed cells (^¥^p<0.05; ^¥¥^p<0.01; ^¥¥¥^p<0.001) were determined using the Student *t*-test.

## DISCUSSION

The present study provides a comprehensive analysis of the roles and limits of BA glucuronidation as a self-defense mechanism developed by liver cells against the accumulation of toxic BAs during chronic or acute cholestatic situations.

Compared to non-cholestatic donors, the serum BA-G profile from BO patients reveals that severe cholestasis affects both the capability and selectivity of bile acid glucuronidation. The 4-fold accumulation of serum BA-G associated with large increases in MRs for certain BA species, including CDCA-3G, CA-24G, LCA-3G, DCA-3G, and DCA-24G, demonstrates that BA glucuronidation is strongly reinforced during acute cholestasis. In these samples, the major components of the BA-G pool correspond to conjugates formed from primary and secondary acids, instead of 6α−hydroxylated BA species, i.e., HDCA and HCA, as in control samples. This reveals that the improved glucuronidation capacity is redirected against the most hepatotoxic BA molecules(4, 5, 11, 29, 41, 43, 44). Indeed, primary and secondary acids exert strong hepatotoxic effects (11, 12, 45), while HDCA and HCA are low toxicity species (11, 12). Results from primary human hepatocytes demonstrate that the mobilization of glucuronidation reflects a feed-forward mechanism by which BAs stimulate their own glucuronidation. Toxic primary and secondary BAs (11, 12, 45) also act as potent activators for the BA sensors FXR (46) and PXR (47). Thus, the herein identification of functional PXREs within the UGT1A3, 1A4, and 2B4 gene promoters, added to the previous findings of FXREs in BA-conjugating UGT gene promoters (26, 27), establish the FXR- and/or PXR-dependent manner in which toxic BAs selectively activate UGT gene expression. The benefits of self-induced BA glucuronidation can be envisioned from the reduced activation of proinflammatory mediators observed in HepG2 cells exposed to BA-G. These BA-G also display reduced pro-apoptotic effects when compared to their unconjugated precursors (11), which indicates that glucuronidation protects hepatic cells against the 2 major hallmarks of BA-induced toxicity, namely the release of pro-inflammatory mediators and the promotion of apoptotic cell death (11, 13, 41, 48). In sum, the present study establishes glucuronidation as a feed-forward mechanism allowing toxic BAs to induce their own glucuronidation, in FXR- and PXR-dependent manners (**Figure 7**). Such a mechanism may reinforce other defense processes developed by liver cells to limit the extent of liver injury caused by BA accumulation (4, 5, 11, 41). These processes include reduced expression of the cytochrome P450 (CYP)7A1 enzyme and Na^+^-taurocholate cotransporting polypeptide (NTCP), i.e., the 2 rate-limiting factors for BA synthesis and uptake, respectively, and activation of hepatic BA-exporting transporters, such as the bile salt export pump (BSEP), the organic solute transporters (OST)s alpha and beta, and the multidrug resistance-associated proteins (MRP) 2, 3 and 4 (*reviewed in* (*15*)). Thus, the mobilization of glucuronidation constitutes an additional “*defensive line*” against BA toxicity in cholestatic liver cells (**Figure 7**).

**Figure 7.**
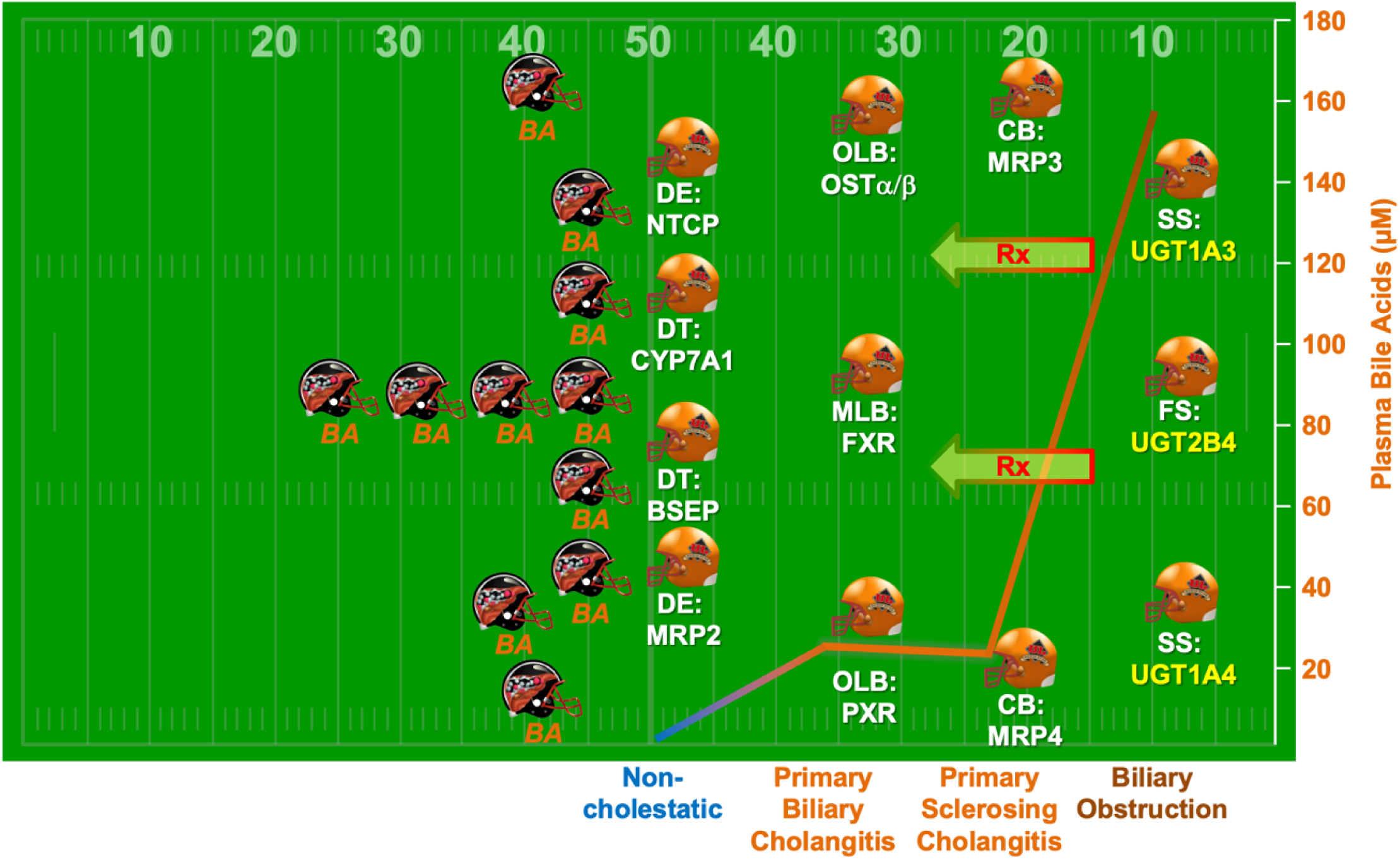
Glucuronidation: the last defensive line against bile acid toxicity during cholestasis. When compared to non-cholestatic donors, patients suffering from primary biliary cholangitis (PBC), primary sclerosing cholangitis (PSC), and biliary obstruction (BO) accumulate toxic bile acids (BA) in their circulation (4, 5, 29). In liver cells, these BAs activate the Farnesoid X- (FXR) and Pregnane X-(PXR) Receptors, which in turn regulate a series of key genes controlling BA synthesis (cytochrome P450, [CYP]7A1), uptake (Na^+^-taurocholate co-transporting polypeptide [NTCP]), apical (multidrug resistance-associated protein [MRP2], and bile salt export pump [BSEP]), and basolateral (MRP3 and 4, organic solute transporters [OST]s alpha and beta) export. Results of the present study identify the FXR- and PXR-dependent manner in which toxic acids activate the hepatic expression of the BA-conjugating UDP-glucuronosyltransferase (UGT)1A3, 1A4, and 2B4. This feed-forward mechanism allowing BAs to self-induce their own glucuronidation is, however, mobilized only in the presence of the huge BA accumulation observed during BO, thus identifying glucuronidation as a “*last defensive line*” against bile acid toxicity. However, it can be envisioned that drugs (*Rx*) targeting UGTs may be efficient in reducing BA toxicity in patients suffering from chronic cholestastic diseases, such as PBC and PSC, like in a “*stop the run defensive way”*. Golden helmets correspond to the official and distinctive uniform of the *Rouge et or*” (red and golden) football team from Laval University (Québec city, Canada), winners of the Canadian University Football championship (Vanier’s cup) in 1999, 2003, 2004, 2006, 2008, 2010, 2012, 2013, 2016, 2018, 2022 and 2024. Positions: DE: defensive end; DT: defensive tackle; MLB: middle linebacker; OLB: outside linebacker; CB: cornerback; SS: strong safety and FS: free safety. Black and golden helmets were created for this publication. Written permission has been obtained from the «*Service des activités sportives de l’Université Laval*» to use the «*Rouge et Or*» logo for this publication.

However, compared to non-cholestatic donors, the relatively minor changes observed in plasma BA-G levels and MRs from PBC and PSC samples indicate that the 9-fold accumulation of BAs detected in these donors (4) is insufficient to efficiently activate glucuronidation. Accordingly, the expression of BA-conjugating UGTs displays only moderated modulation in hepatocytes exposed to BA mixtures (4) mimicking the PBC and PSC situations. Furthermore, the limited activation of BA glucuronidation during cholestasis is also supported by the absence of UGT2B and UGT1A protein accumulation in PBC and PSC liver samples. This last observation is consistent with previous reports indicating that UGT2B4 and 2B7 mRNA levels were only mildly altered in PBC livers (49). These results therefore indicate that, in contrast to biliary obstruction, PBC and PSC situations do not lead to significant mobilization of BA glucuronidation. By contrast, other BA detoxification systems, namely the induction of OSTα/β (50), MRP3 (51, 52), MRP4 (49) levels and repression of CYP7A1 (49), NTCP (51), and MRP2 (53) expression are already activated in PBC and/or PSC livers (**Figure 7**). In sum, the differential response of UGT expression during acute, i.e., BO, *versus* chronic, i.e., PBC and PSC, cholestasis identifies glucuronidation as a “*last defensive line*” against BA toxicity (**Figure 7**), which is mobilized only under severe cholestatic situations.

However, even if *in vitro* experiments illustrate an impressive reduction of BA toxicity after glucuronidation, the benefits of this “*last defensive line*” may be limited in clinics even in patients with acute cholestasis. Indeed, the differential accumulation between non-glucuronidated (58-fold) (5) and glucuronidated (4-fold) bile acids in BO sera indicates that when toxic acids activate their own glucuronidation, they already have reached such elevated concentrations that improved glucuronidation may only have moderate impacts on the total BA toxicity. Thus, it appears that this endogenous feed-forward mechanism is a “*too little, too late*” defensive reaction. Nevertheless, the present work provides experimental and clinical rationales to further consider glucuronidation as a potential pharmacological option for filling the vacuum in the PBC and PSC therapeutic armamentarium.

Until recently, the therapeutic options for PBC patients were limited to UDCA (Ursodiol®), (OCA, Ocaliva®), and ultimately liver transplantation (1, 2, 54). However, up to 40% of PBC patients do not respond satisfactorily to UDCA, and no medical therapies have been proven to delay PSC progression (1, 3, 54–56), thus illustrating the need for alternative pharmacological options. Considering that a reduction of BA toxicity in these cholestatic pathologies remains a major therapeutic goal (14), pharmacological activation of hepatic BA-conjugating UGT expression may further reinforce the hepatoprotective properties of glucuronidation and thus correct the “*too little*” nature of its activation (**Figure 7**). Interestingly, the European, American and Asian medical associations also recommend the use of PPAR activator such as Fenofibrate and Bezafibrate as second-line treatment option for PBC. Interestingly, fenofibrate which reduces ALP levels in PBC and PSC patients, also induced serum BA-G levels in PBC and PSC patients (25). Furthermore, additional analyses identified serum BA-glucuronides as measures of response to fenofibrate therapy in these patients (25), thus supporting the idea that increased formation of BA-G may be associated with improved biochemical outcomes in PBC and PSC patients. In 2024, 2 novel PPAR agonists, namely Elafibranor (Irqvo®) (57) and Seladelpar (Livdelzi®) (58), received FDA approval for this disease PBC. Investigating how they impact BA glucuronide formation would be of interest.

In conclusion, the present study identifies a feed-forward mechanism allowing BAs to self-stimulate their own glucuronidation in PXR- and/or FXR-dependent manners during acute cholestasis. The “*too little, too late*” nature in which this self-defense mechanism is mobilized during chronic and acute cholestasis may limit its hepato-protective properties, but the reduced toxicity exhibited by BA-G strongly encourages the search for novel therapeutics acting as inducers of BA-glucuronidation UGTs (**Figure 7**). Even if additional investigations with larger studies are required to further evaluate how BA glucuronide conjugation evolves under the different stages of cholestasis, these data establish glucuronidation as a potential pharmacological target in the treatment of cholestatic diseases.

## Abbreviations

ALT: alanine aminotransferase
ALP: alkaline phosphatase
AST: aspartate aminotransferase
BA: bile acid
BO: biliary obstruction
BSEP: bile salt export pump
CA: cholic acid
CDCA: chenodeoxycholic acid
CYP: cytochrome P450
DCA: deoxycholic acid
DR: direct repeat
EMSA: electrophoretic mobility shift assay
ER: everted repeat
FXR: farnesoid X-receptor
FXRE: FXR response element
G: glucuronide
HDCA: hyodeoxycholic acid
HCA: hyocholic acid
ICAM: intercellular adhesion molecule
IL: interleukin
LCA: lithocholic acid
LC-MS/MS: liquid chromatography-tandem mass spectrometry
MRP: multidrug resistance-associated proteins
NTCP: Na^+^-taurocholate cotransporting polypeptide
OST: organic solute transporter
PAI: plasminogen activator inhibitor
PBC: primary biliary cholangitis
PPAR: peroxisome proliferator activated receptor
PSC: primary sclerosing cholangitis
PXR: pregnane X-receptor
PXRE: PXR response element
UDCA: ursodeoxycholic acid
UGT: UDP-glucuronosyltransferase.

## ACKNOWLEDGEMENT

We wish to thank Dr. Virginie Bocher for critical reading of the manuscript, and Professor Jenny Heathcote from the Toronto Health Network for providing clincal samples.

## AUTHOR’S CONTRIBUTIONS

Jocelyn Trottier, Audrey-Anne Lavoie participated to the study design, acquisition and analysis of data and manuscript drafting and revision. Ewa Wunsch and Mélanie Verreault participated to the acquisition, analysis and interpretation of data as well as to the manuscript drafting and revision. Olivier Barbier, Robert J. Straka, Jenny Heathcote, Nisanne Ghonem and Piotr Milkiewicz were involved in the study concept and design, analysis and interpretation of data, and manuscript drafting and revision.

## DATA AVAILABILITY STATEMENT

Data available on request from the authors.

## DECLARATION OF GENERATIVE AI AND AI-ASSISTED TECHNOLOGIES IN THE WRITING PROCESS

During the preparation of this work the authors did not use any generative AI tool.

## Supplemental material 1: Additional experimental procedures

### Materials

Chenodeoxycholic acid-3glucuronide (CDCA-3G), CDCA-24G, deoxycholic acid-3G (DCA-3G), DCA-24G, hyodeoxycholic acid-6G (HDCA-6G), HDCA-24G, lithocholic acid-3G (LCA-3G), LCA-24G, hyocholic acid-24G (HCA-24G) and cholic acid-24G (CA-24G) were synthesized by the organic synthesis service at the CHU de Québec Research Centre (http://pfchem.crchul.ulaval.ca/en/index.html) (Québec, Canada)(21). Deuterated [^2^H_4_]-CDCA, [^2^H_4_]-DCA, [^2^H_4_]-LCA, and [^2^H_4_]- CA were purchased from C/D/N Isotopes (Montréal, Canada). The preparation of [^2^H_4_]-CDCA-24G, [^2^H_4_]-DCA-3G, [^2^H_4_]-DCA-24G, [^2^H_4_]-LCA-24G and [^2^H_4_]-CA-24G was as reported(59). Unconjugated, taurine- and glycine-conjugated bile acids were purchased from Steraloids (Newport, RI). Methanol and isopropanol were purchased from VWR (Montréal, Canada). Ammonium formate was obtained from Laboratoire Mat (Québec, Canada). All the reagents were of the highest grade commercially available. Strata X and Synergi Hydro-RP columns were obtained from Phenomenex (Torrance, CA). Cryopreserved human hepatocytes from 3 individual donors (SM7) were obtained from Celsis-InVitro Technologies (Baltimore, MD), and cultured as reported previously(23, 36). Human hepatoma HepG2 cells were from the American Type Culture Collection (Rockville, MD) and were grown as described(23, 36). Cell culture reagents (including the Exgen500 transfection reagent) were from Fermentas (Ottawa, Canada). GW4064 and Rifampicin were from Sigma (St. Louis, MO), and the SYBR Fast PCR mix was purchased from Life Technologies (Foster City, CA). The Tri-Reagent acid: phenol reagent was from Bioshop (Burlington, Canada). Protein assay reagents were obtained from Bio-Rad Laboratories Inc. (Marnes-la-Coquette, France). The anti-UGT1A and anti-UGT2B antibodies(16, 26) were kindly provided by Dr. A. Bélanger (Laval University, Québec, Canada). The rabbit polyclonal anti-human actin antibody was purchased from Sigma (A5060), and the secondary anti-rabbit IgG HRP-linked (NA934) was obtained from GE Healthcare (Burnaby, Canada). Hyperfilm were from Amersham (Oakville, Canada). The anti-PXR antibody (sc-9690) was from Santa Cruz Biotechnology (Santa Cruz, CA). γ-^32^P-ATP was purchased from NEN-Life Sciences (Boston, MA). Quick Change Site Directed Mutagenesis Kit was purchased from Stratagene (La Jolla, CA).

### Blood donors

Liver biochemistries from non-cholestatic volunteers (controls) and cholestatic patients involved in serum BA-G analyses are shown in SM3.

### Non-cholestatic blood donors

The non-cholestatic donors (control population) for this study were selected among the participants of the Genetics of Lipid Lowering Drugs and Diet Network (GOLDN) study, a single-arm, uncontrolled, nonrandomized intervention aimed at identifying genetic factors associated with interindividual variability of the triglyceride response to high-fat meals and fenofibrate(21, 29–31). Only participants who had not taken lipid-lowering agents for at least 4 weeks before the initial visit were included. Exclusion criteria were as reported(30), and included elevated lipids and serum concentrations of liver enzymes (aspartate aminotransferase and alanine aminotransferase)(30). As extensively described in Lai et al.(30), participants took part in five visits. However, sera used in the present study were drawn at the first visit, when patients were free of any drug treatment or lipid challenge.

### BO, PBC, and PSC blood donors

Seventeen biliary obstructed (BO) patients (8♂, 9♀) with clinical and biochemical features of cholestasis were recruited(5). In all cases, dilatation of the biliary tree was first confirmed with abdominal ultrasound. Diagnoses were: common bile duct stones: 8; pancreatic tumor: 6; bile duct tumor: 1; benign common bile duct stenosis: 1; and chronic pancreatitis: 1. The endoscopic retrograde cholangiopancreatography (ERCP) procedure involved placement of plastic 10F stents which were removed or replaced at a median of 5 weeks after the initial procedure, depending on the diagnosis.

Eight patients with PBC (PBC-Canada, all women) were recruited at the Toronto Western Hospital (University Health Network, Toronto, Canada)(4). This population comprised 1 patient at stage 0, 3 at stage 1, 3 at stage 2, and 1 cirrhotic patient at stage 4. A second set of 4 PBC patients (PBC-Poland: all women) were recruited at the Pomeranian Medical University (Szczecin, Poland)(4). Three of them were at stage 1, and 1 was cirrhotic. None of the BAs analyzed displayed a significant difference between the two PBC groups (SM2), and results from these 2 PBC sub-populations were mixed for subsequent analyses.

Six patients with PSC were also recruited at the Pomeranian Medical School. Only 1 out of them showed features of liver cirrhosis on imaging studies (abdominal ultrasound, CT-scan, magnetic resonance cholangiopancreatography (MRCP) and/or endoscopic cholangiography (ERCP)(4). In addition, 4 PSC patients also had ulcerative colitis. MRCP/ECRP procedures did not allow the stratification of these patients within the disease stages and liver biopsy is not required for establishing the diagnosis of PSC. The blood samples used in this study were drawn shortly after establishing the clinical diagnosis. None of the PBC or PSC patients was receiving Ursodiol® at the time of blood sampling(4).

In all cases, blood was drawn after a 12H fast, and serum was purified and frozen at -80 °C until analyses.

### Non-cholestatic, PBC and PSC liver donors

Non-cholestatic liver samples were a kind gift from Prof. Ted T. Inaba (Department of Pharmacology, University of Toronto, Toronto, Canada)(33). Tissues were from kidney donors of Caucasian origin (SM4) and were obtained at the time of kidney donation. Tissues were quickly immersed in 1.15%KCl at 4°C until they could be snap-frozen in liquid nitrogen, and were stored at -80 °C at the University of Toronto, Department of Pharmacology, as previously described(33).

The study group consisted of 22 cirrhotic patients. Detailed demographic data of PBC and PSC donors for liver tissue samples are shown in SM5. Liver tissue samples were collected from explanted livers during liver transplantation and immediately frozen at -80 ° C. All patients with PBC and PSC were treated with Ursodiol® at the time of the tissue sampling. The diagnosis of PSC donors (8 men and 2 women) was confirmed by imaging techniques (MRCP or ERCP) and/or liver histology. The diagnosis of PBC donors was according to the widely accepted criteria, which included: elevated ALP, typical liver histology and positive titers of antimitochondrial autoantibodies (AMA). Liver cirrhosis was confirmed with liver biopsy or typical appearance of the liver on abdominal ultrasound and/or computed tomography (CT) scan.

### Cell culture and mRNA levels determination

For RNA and protein analyses, 350,000 hepatocytes/well were seeded in 12-well plates and cultured in the *Invitro Gro CP* medium for 48H. Total RNA was isolated from treated or control cells according to the Tri-Reagent acid: phenol protocol, as specified by the supplier (Bioshop). The reverse transcription reaction was performed using 200 units of Superscript II (Invitrogen) with 1µg of total RNA, and 7.5ng of random hexamer (Roche, Laval, Canada) at 42 °C for 50 min as described(23, 36). The quantitative RT-PCR reactions were performed with gene-specific primers as listed in SM9, using an ABI Prism 7500FAST instrument from Applied Biosystems. For each reaction, the final volume of 10 µL was comprised of 5 µL of SYBR Fast PCR Mix, 1 µL of each primer (4 µM), and 3 µL of diluted RT products. Conditions for real-time PCR were 95°C for 20sec, 95°C for 1sec and annealing temperature (SM9) for 20 sec for 40 cycles(21). For each gene and each cell line, the amplification efficiency and the accuracies of ΔΔCt versus the housekeeping 28S (human hepatocytes)(21) or PUM-1 (HepG2 cells)(21) genes were tested using 2 to 5 log of concentrations of cDNA produced from Hepatocytes- or HepG2-cell purified mRNA. C_T_ values were analyzed using the comparative C_T_ (ΔΔC_T_) method as recommended by the manufacturer (Life Technologies). The amount of target (2^−ΔΔCT^) was obtained by normalizing to the endogenous reference 28S or PUM-1 and was relativized to the vehicle-treated baseline control(21).

### Glucuronidation assays

Control- and rifampicin-treated HH were homogenized in PBS-dithiothreitol (0.5 mM). All assays were performed for 1H as previously reported(23, 60) using 10 0µM BAs, 1 mM UDPGA, and 5-10 µg homogenates. BA-G formation was quantified by LC-MS/MS.

### Microsomal protein isolation and western-blot analyses

Microsome pellets from non-cholestatic, PBC, and PSC livers were purified and resuspended at 5 µg/µL, as previously reported(23). For western-blot analyses, liver microsomes (10 µg) or total proteins from hepatocytes (10 µg) were size-separated by 10% SDS-PAGE, transferred onto nitrocellulose membranes, and then hybridized with the anti-UGT1A (1:2,000) or anti-UGT2B (1:2,000) antibodies, as previously described(21, 23). An anti-rabbit IgG horse antibody (1:10,000) conjugated with peroxidase was used as the second antibody. Immunocomplexes were visualized on hyperfilm. When appropriated, the same membranes were then rehybridized with an anti-actin (1:2,000) antibody as a loading control assessment.

### Electrophoretic mobility shift assay (EMSA)

EMSA using *in vitro* produced PXR and RXR(36) were performed using the radiolabeled probes listed in SM9. Annealed oligonucleotides were end-labeled with γ-^32^P-ATP using T4-polynucleotide kinase (Roche), while the PXR and/or RXR proteins were synthesized *in vitro* using the TNT Quick Coupled Transcription/Translation System (Promega, Madison, WN). Proteins (2 µL) were incubated for 15 min at 4 °C in a total volume of 20 µL of EMSA buffer (10 mM Hepes, pH 7.8; 60 mM KCL; 0.2% Igepal; 6% Glycerol; 2 mM DTT; 2.0 µg dIdC and 1 µg SS-DNA). Subsequently, 100,000 cpm of double-stranded and radiolabeled oligonucleotides were added and further incubated for 15 min at room temperature. The protein complexes were resolved by 6% non-denaturing polyacrylamide gel electrophoresis. For competition experiments, unlabeled oligonucleotides were included in the binding reaction at 1-, 10-or 50-fold excess concentrations over the probe. For supershift experiments, the anti-PXR antibody (Santa-Cruz, sc-9690) was pre-incubated for 20 min in the binding buffer before the addition of PXR and RXR proteins.

### Plasmid cloning, site directed mutagenesis and transient transfection assays

Luciferase reporter plasmids for the wild type (WT) or mutated (MT), UGT1A3 and/or 1A4 PXR response elements (PXREs) were obtained and transiently transfected into HepG2 cells by cloning 3 copies of the dimerized oligonucleotides (SM9) in the thymidine kinase promoter-driven luciferase reporter (TKpGL3) vector following the previously reported procedure(24). The WT UGT2B4p-524 reporter construct was described previously(24), and mutations in the PXRE from this construct were introduced using the Quick Change Site Directed Mutagenesis Kit (Stratagene) and the oligonucleotide UGT2B4 DR4-490 bpmt (SM9). HepG2 cells were transfected with 100 ng of the indicated luciferase reporter plasmids, 10 ng of the pRL-NULL expression vector, and 30 ng of PXR and RXR expression plasmids(24, 34) using Exgen500 (Fermentas). After 6H at 37 °C for transfection, cells were incubated for 24H with either DMSO (vehicle) or rifampicin (20 µM). Luciferase and renilla activities were determined as reported(24).

## Supplemental material 2. Comparison of the bile acid-glucuronide profiles in the 2 PBC populations.

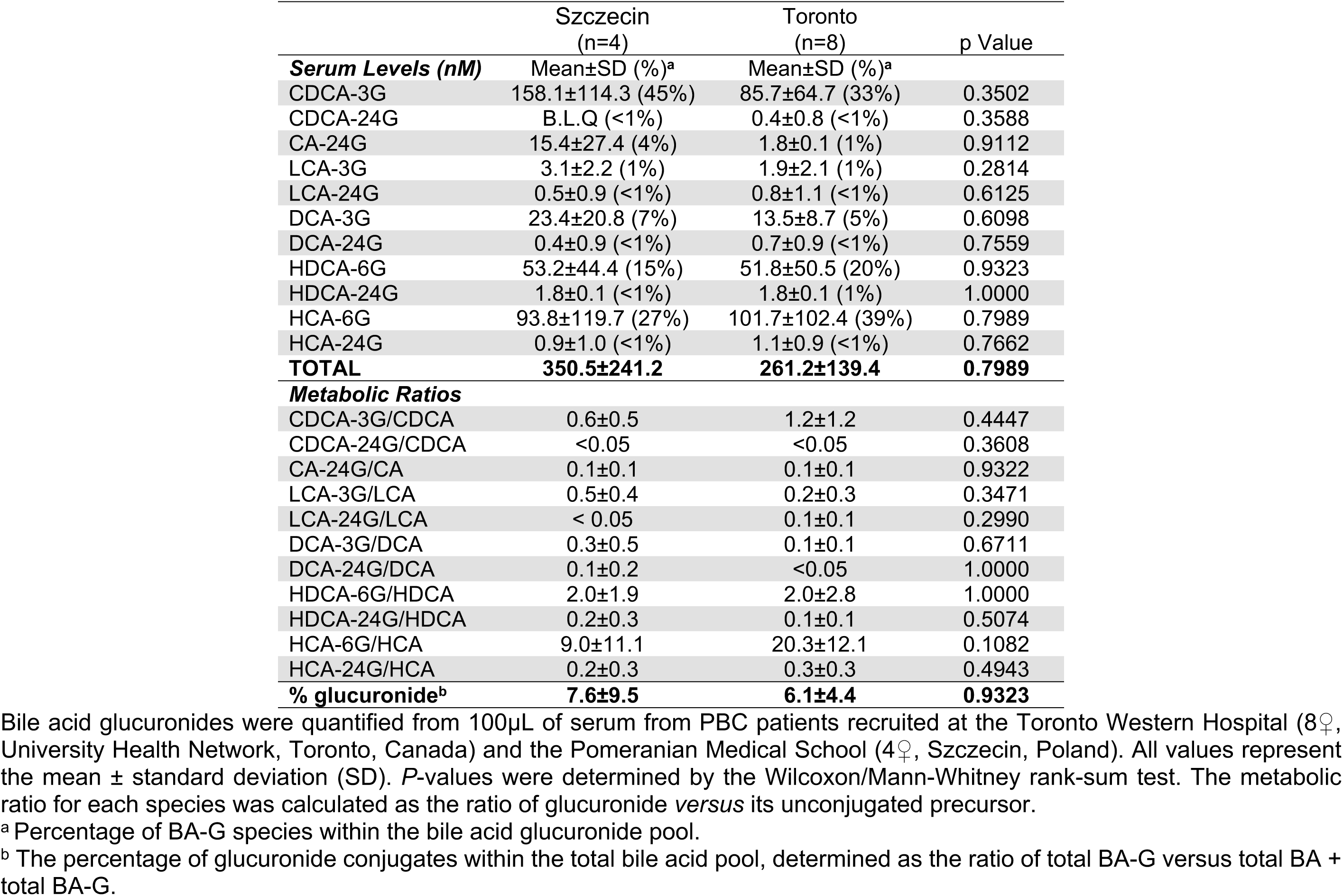
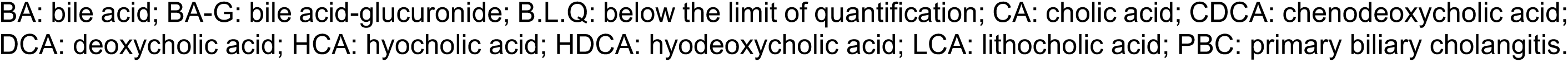

## Supplemental material 3. Demographic and laboratory characteristics of donors for serum samples.

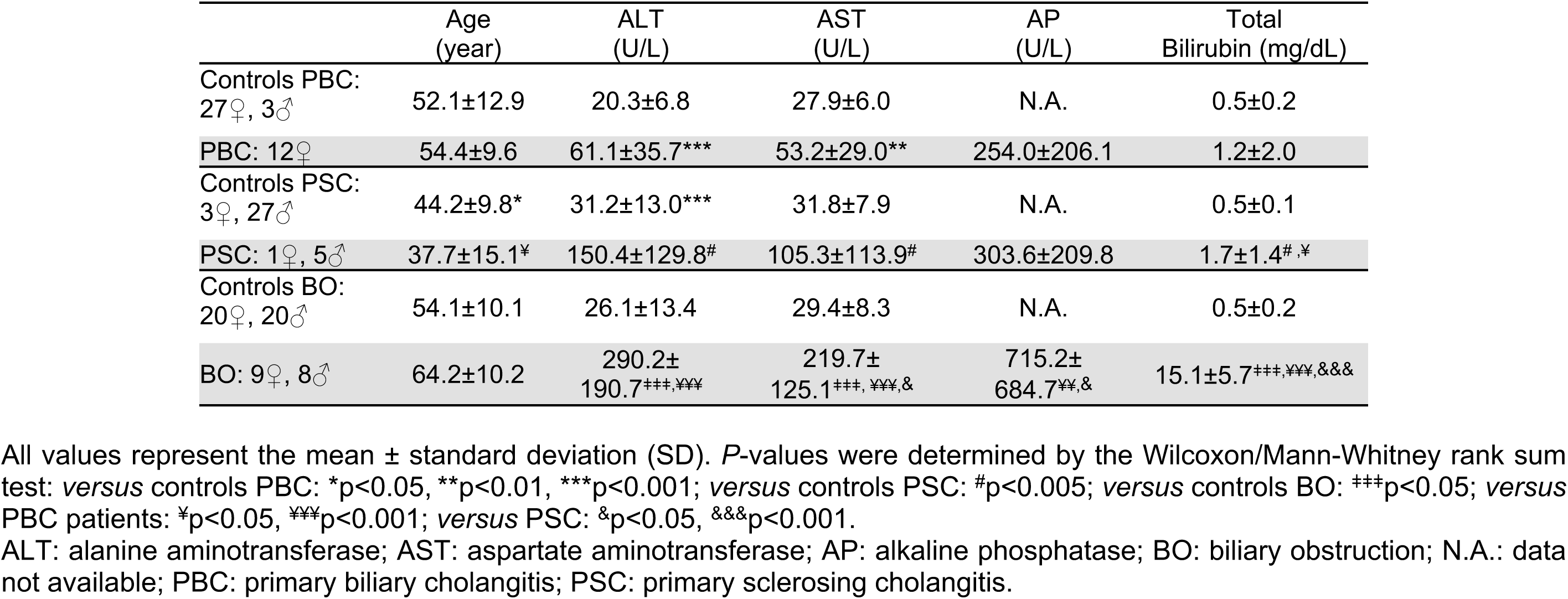

## Supplemental material 4. Demographic and laboratory characteristics of non-cholestatic liver donors.

Derived from Sumida *et al*(*33*).

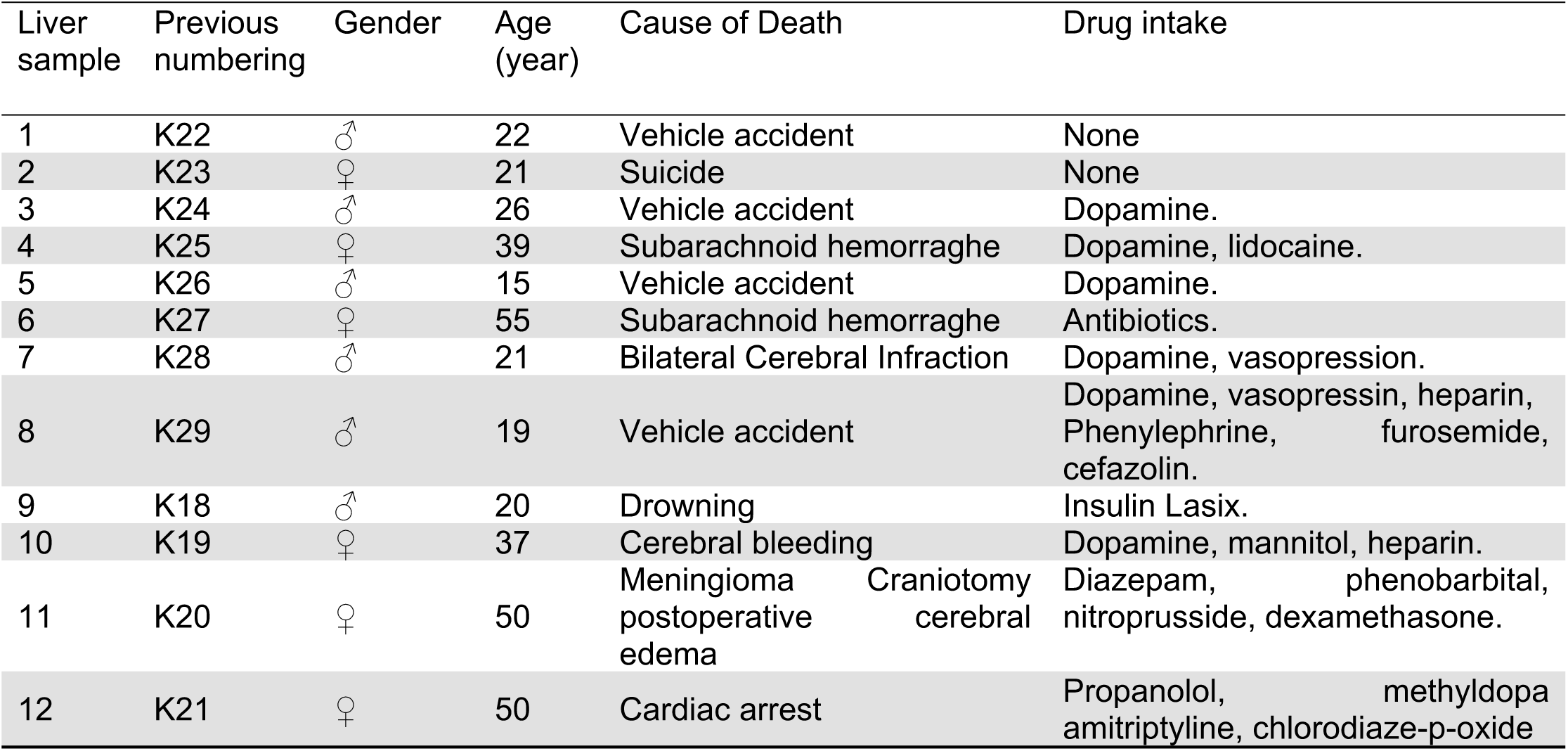

## Supplemental material 5. Demographic and laboratory characteristics of PBC and PSC liver transplant recipients whose explanted livers were used in the study.

|  | PBC<br>10♀, 2♂ | PSC<br>2♀, 8♂ |
| --- | --- | --- |
| Age (year) | 55.5±8.6 | 46.9±12.6 |
| AST(U/L) | 178.7±135.6 | 516.0±585.0 |
| ALP (U/L) | 628.0±392.3 | 623.2±351.3 |
| Total Bilirubin (mg/dL) | 6.7±6.8 | 6.3±5.3 |
| Albumin (g/L) | 25.0±6.1 | 24.2±5.8 |
| Creatinine (μmol/L) | 109.4±106.0 | 79.9±11.4 |

All values represent mean ± standard deviation (SD). AP: alkaline phosphatase; AST: Aspartate aminotransferase; PBC: primary biliary cholangitis; PSC: primary sclerosing cholangitis.

## Supplemental material 6. Comparison of serum bile acid-glucuronide profiles within the 3 control groups.

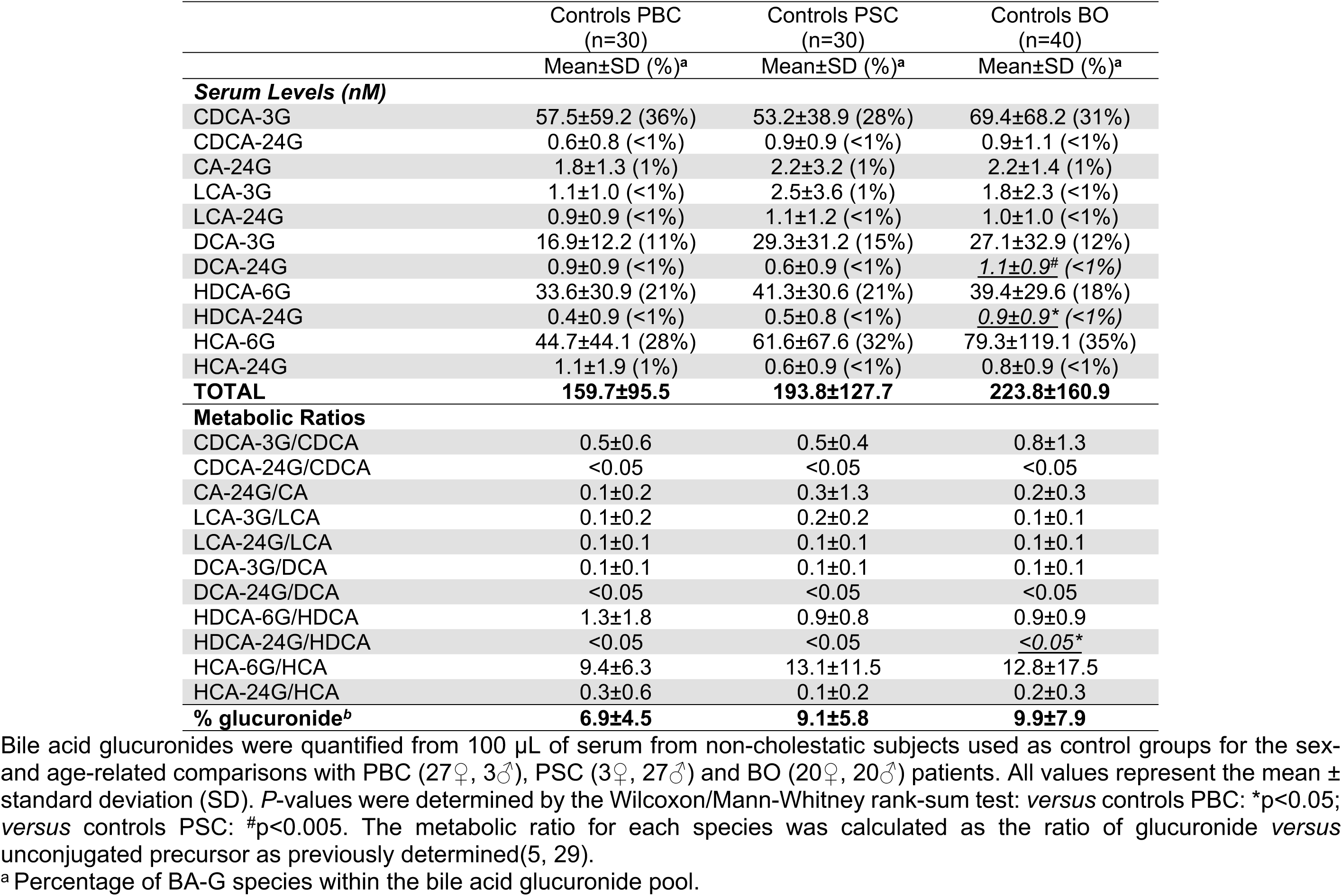
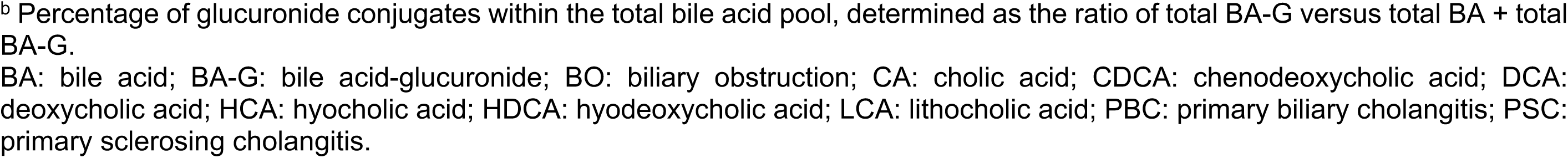

## Supplemental material 7. Demographics and causes of death of donors for human hepatocytes.

| Donor | Previous numbering | Treatment/ Analysis | Gender | Age (year) | Race | Cause of Death |
| --- | --- | --- | --- | --- | --- | --- |
| 1 | CPQ | BA & GW4064/ mRNA&protein | ♂ | 31 | Caucasian | Head trauma 2<sup>nd</sup> to MVA |
| 2 | TQS | Rifampicin/ mRNA&protein | ♂ | 46 | Caucasian | Stroke |
| 3 | ZYZ | Rifampicin/ mRNA&protein | ♂ | 46 | Afr.-Amer. | Anoxia |
| 4 | AAA | Rifampicin/ Glucuronidation | ♀ | 8* | Afr.-Amer. | Head Trauma |
| 5 | BAD | Rifampicin/ Glucuronidation | ♂ | 60 | Caucasian | Intracerebral hemorrhage |

Afr.-Amer.: African-American; BA: bile acids (SM8); MVA: Malignant ventricular arrhythmia
\*age: weeks

## Supplemental material 8. Composition of the bile acid mixtures used with human hepatocytes in order to mimic the profiles detected in sera from the non-cholestatic volunteers, PBC, PSC and BO patients as previously reported(4, 5, 29).

|  | Non-cholestatic | PBC | PSC | BO |
| --- | --- | --- | --- | --- |
| CDCA | 257.0 | 217.0 | 152.0 | 85.0 |
| TCDCA | 120.0 | 4,100.0 | 1,400.0 | 16,500.0 |
| GCDCA | 772.0 | 6,300.0 | 7,000.0 | 28,500.0 |
| CA | 182.0 | 122.0 | 110.0 | 173.0 |
| TCA | 180.0 | 8,400.0 | 3,800.0 | 54,500.0 |
| GCA | 233.0 | 4,500.0 | 9,100.0 | 56,200.0 |
| UDCA | 138.0 | 114.0 | 683.0 | 20.0 |
| TUDCA | 5.0 | 27.0 | 35.0 | 75.0 |
| LCA | 13.0 | 15.0 | 6.0 | 2.0 |
| TLCA | 23.0 | 8.0 | 5.0 | 12.0 |
| GLCA | 16.0 | 33.0 | 10.0 | 11.0 |
| LCA-S | 7.0 | 8.0 | 1.0 | 4.0 |
| DCA | 387.0 | 258.0 | 75.0 | 21.0 |
| TDCA | 45.0 | 164.0 | 28.0 | 213.0 |
| GDCA | 246.0 | 912.0 | 93.0 | 418.0 |
| HDCA | 41.0 | 38.0 | 12.0 | 5.0 |
| HCA | 5.0 | 7.0 | 15.0 | 14.0 |
| **TOTAL** | **2,670.0** | **25,223.0** | **22,525.0** | **156,753.0** |

Bile acid concentrations are in nM. CA: cholic acid; CDCA: chenodeoxycholic acid; DCA: deoxycholic acid; HCA: hyocholic acid; HDCA: hyodeoxycholic acid; LCA: lithocholic acid; UDCA: ursodeoxycholic acid; T: tauro-; G: glyco-; -S: sulfate; PBC: primary biliary cholangitis; PSC: primary sclerosing cholangitis; BO: biliary obstruction.

## Supplemental material 9. Oligonucleotides used in this study.

Oligonucleotides were used for real-time PCR analyses, electrophoretic mobility shift assays, site-directed mutagenesis of the previously reported UGT2B4p-524 promoter construct(24) or construction of multiple copies of potential PXR response elements from the UGT1A3, 1A4 and 2B4 genes.

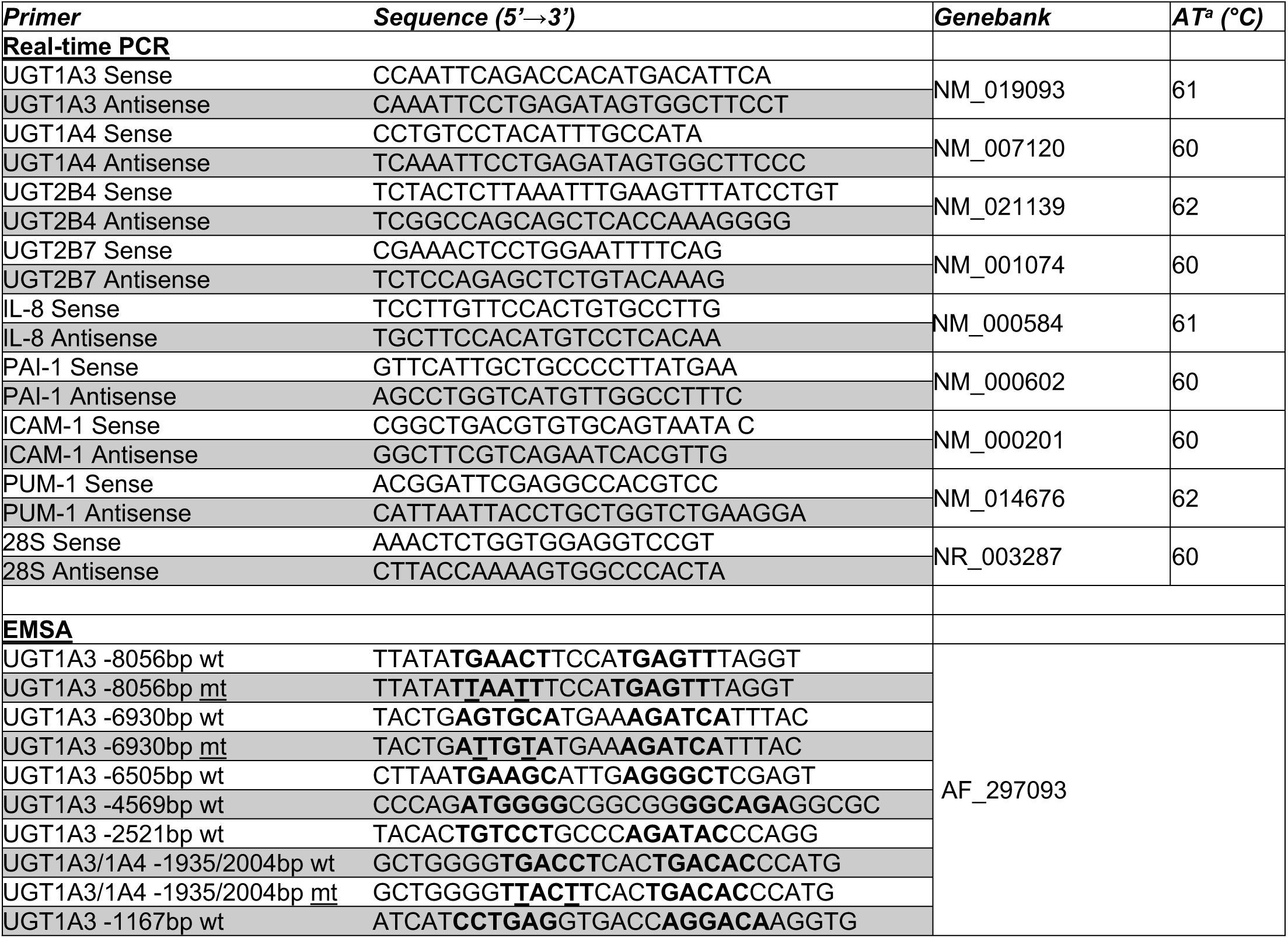
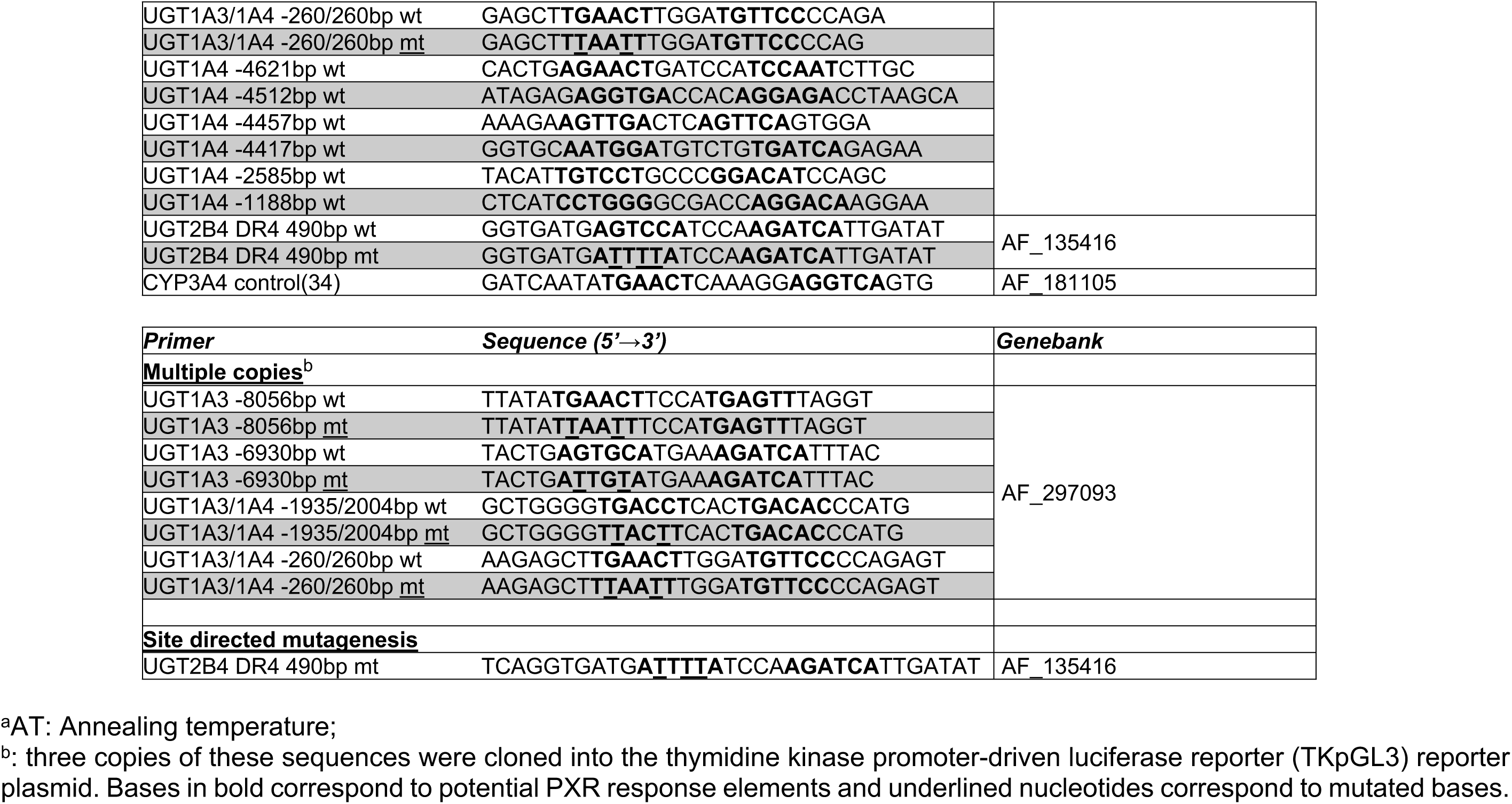

## Supplemental material 10. Changes of the serum bile acid-glucuronide profiles between non-cholestatic controls and patients with Primary Biliary Cholangitis (PBC).

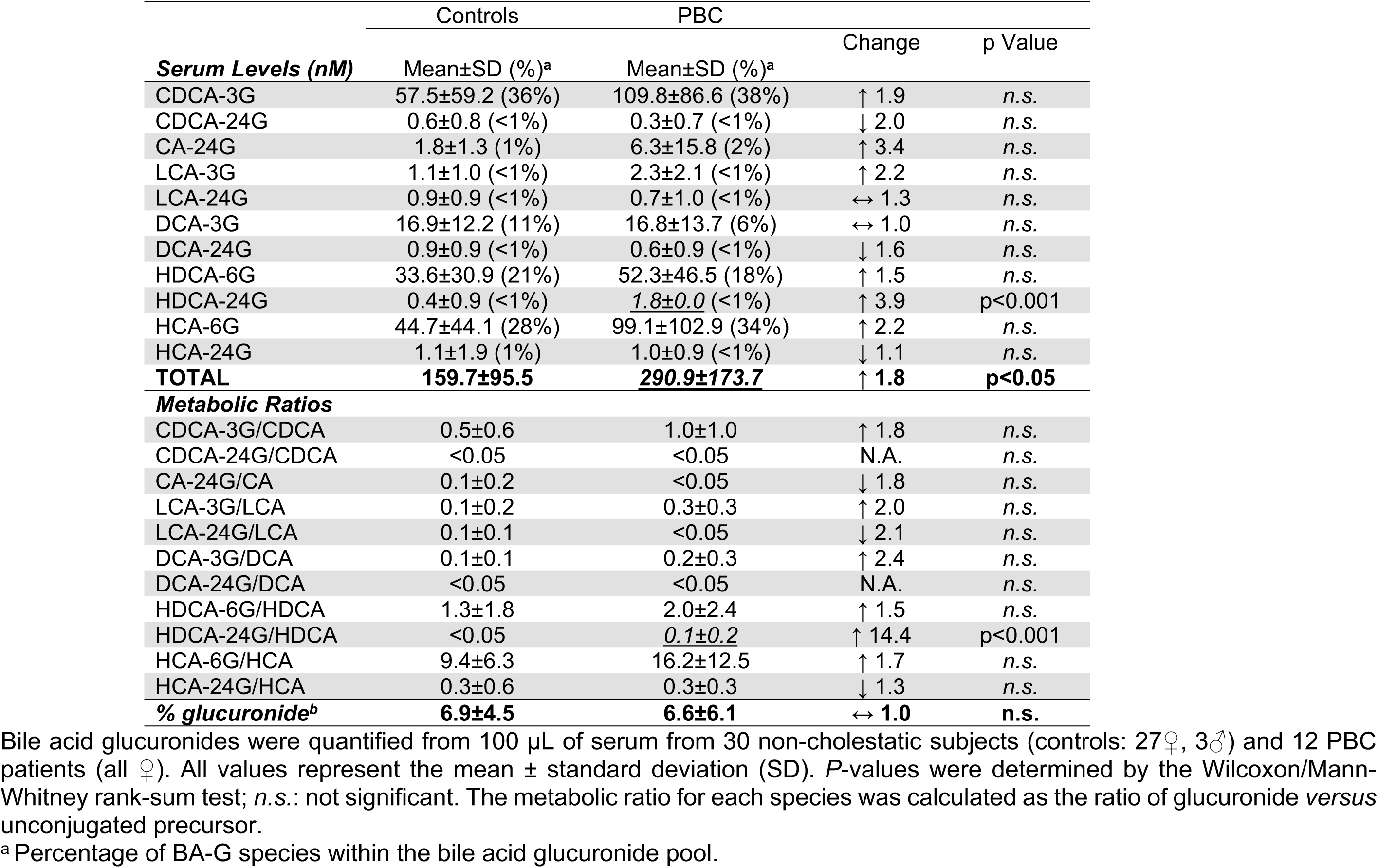
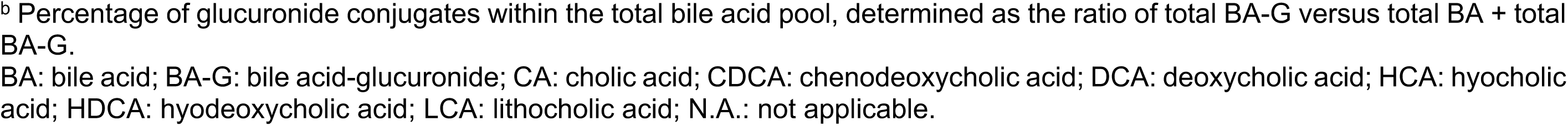

## Supplemental material 11. Changes of the serum bile acid-glucuronide profiles between non-cholestatic controls and patients with Primary Sclerosing Cholangitis (PSC).

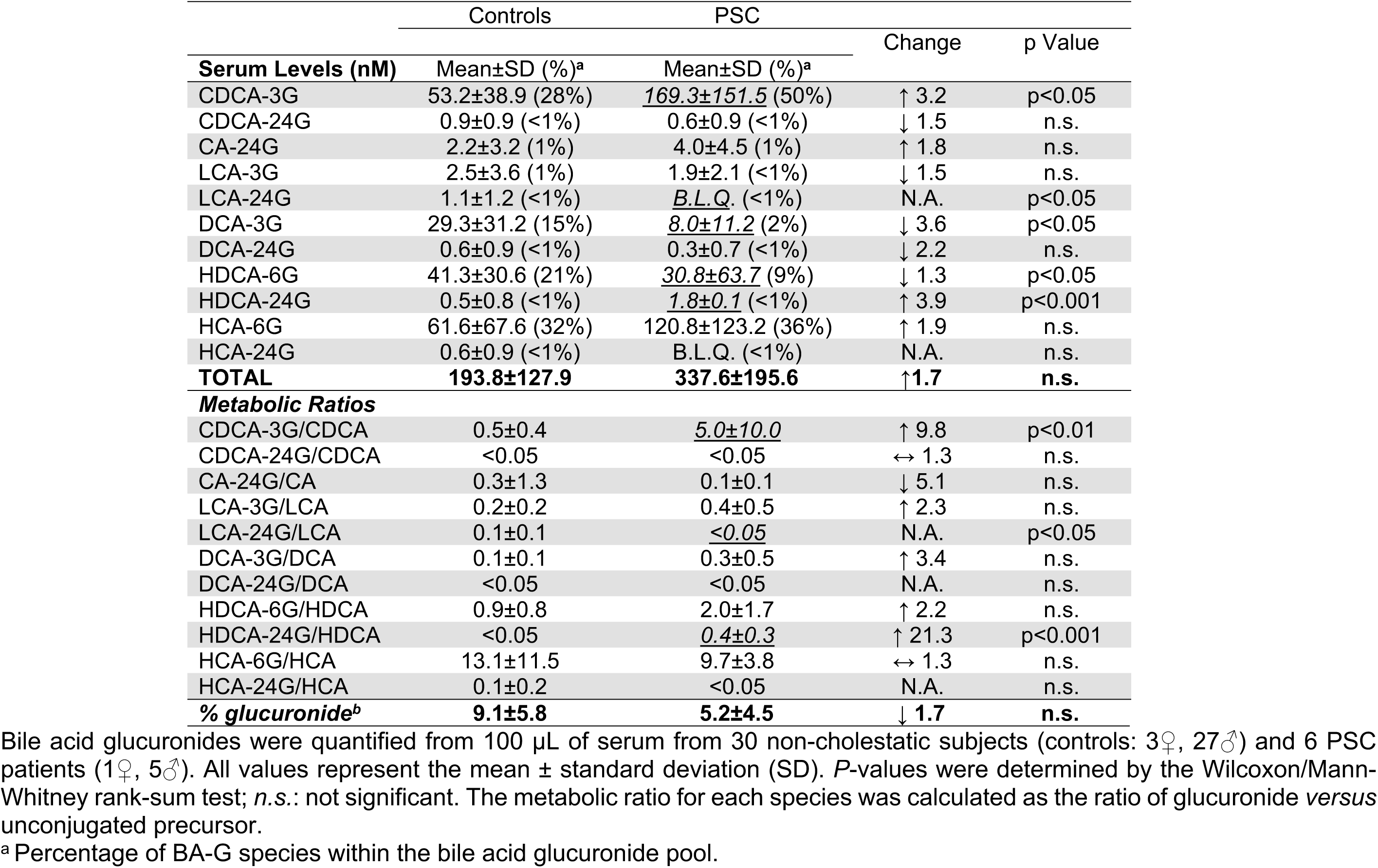
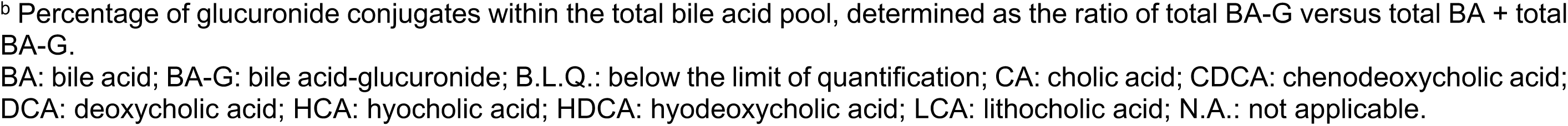

## Supplemental material 12. Changes of the serum bile acid-glucuronide profiles between non-cholestatic controls and patients with Biliary Obstruction (BO).

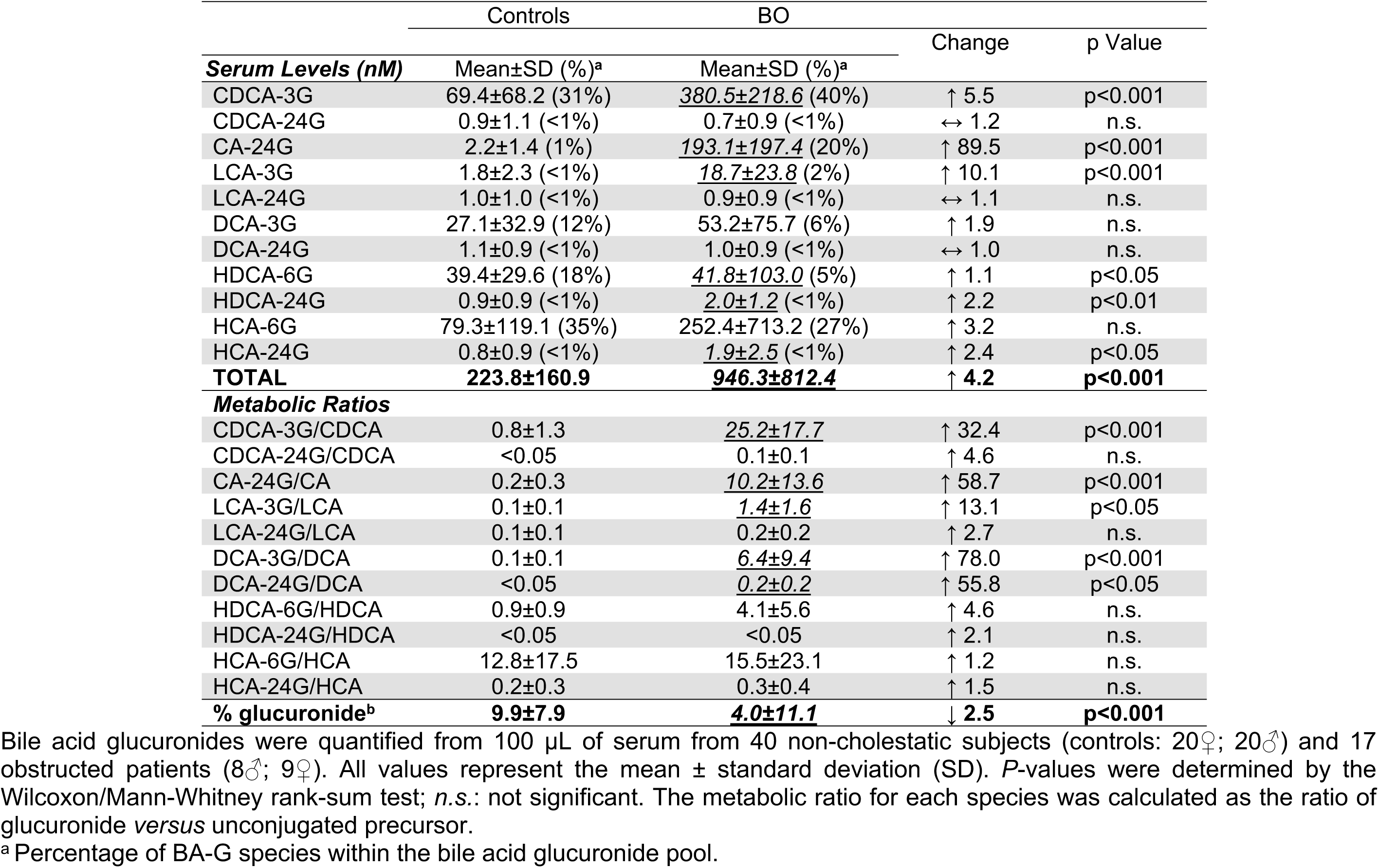
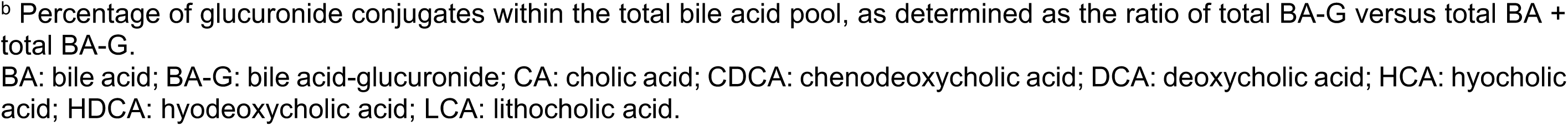

## Supplemental material 13. Rifampicin selectively activates bile acid glucuronidation in human hepatocytes.

| Glucuronide formation | Donor 4 |  |  | Donor 5 |  |  |
| --- | --- | --- | --- | --- | --- | --- |
|  | Vehicle | Rifampicin | *P value* | Vehicle | Rifampicin | *P value* |
| CDCA-3G | 0.23±0.01 | 0.32±0.07 | *0.115* | 0.33±0.02 | 0.44±0.03 | *0.005* |
| CDCA-24G | 10.41±1.17 | 13.59±1.43 | *0.040* | 10.93±0.34 | 13.42±1.82 | *0.079* |
| CA-24G | 0.29±0.02 | 0.37±0.04 | *0.037* | 0.33±0.05 | 0.44±0.04 | *0.097* |
| LCA-3G | 1.47±0.10 | 1.77±0.01 | *0.007* | 1.47±0.14 | 2.45±0.20 | *0.023* |
| LCA-24G | 4.60±0.03 | 5.80±0.24 | *0.001* | 4.55±0.12 | 6.00±0.20 | *0.000* |
| HDCA-6G | 225.96±4.24 | 254.08±14.86 | *0.014* | 265.60±5.95 | 311.03±2.56 | *0.000* |
| HDCA-24G | 59.29±2.56 | 73.60±6.12 | *0.020* | 91.89±7.94 | 127.86±8.61 | *0.006* |
| HCA-6G | 5.82±0.10 | 6.21±0.27 | *0.068* | 6.92±2.59 | 8.23±0.87 | *0.450* |
| HCA-24G | 0.76±0.17 | 0.97±0.06 | *0.064* | 0.52±0.22 | 1.28±0.16 | *0.003* |

Primary human hepatocytes from donors 4 and 5 (SM7) were treated for 48H with DMSO (vehicle) or 20 µM Rifampicin (donors 2-5). Cells were then homogenized in PBS-dithiothreitol (0.5 mM), and assayed for bile acid glucuronidation through 1H *in vitro* assays, performed with 5 to 10 µg homogenates in a previously reported enzymatic buffer (23, 60) enriched with each acid (100 µM) and the co-substrate UDPGA (1 mM). Glucuronide formation was resolved through LC-MS/MS. Values (mean±SD) are expressed as nmol/H/mg proteins. Statistically significant *differences* between control and treated cells are indicated by asterisks (Student *t*-test: \*: p<0.05; \*\*: p<0.01; \*\*\*: p<0.001).
-G: glucuronide; CA: cholic acid; CDCA: chenodeoxycholic acid; HCA: hyocholic acid; HDCA: hyodeoxycholic acid; LCA: lithocholic acid; UDPGA: UDPGA-glucuronic acid.

## Supplemental material 14. Nucleotides sequences of the human UGT1A3 and 1A4 promoter genes derived from GenBank AF_297093, and position of the direct repeat (DR)3, DR4 or everted repeat (ER6) motifs identified as potential PXR response elements.

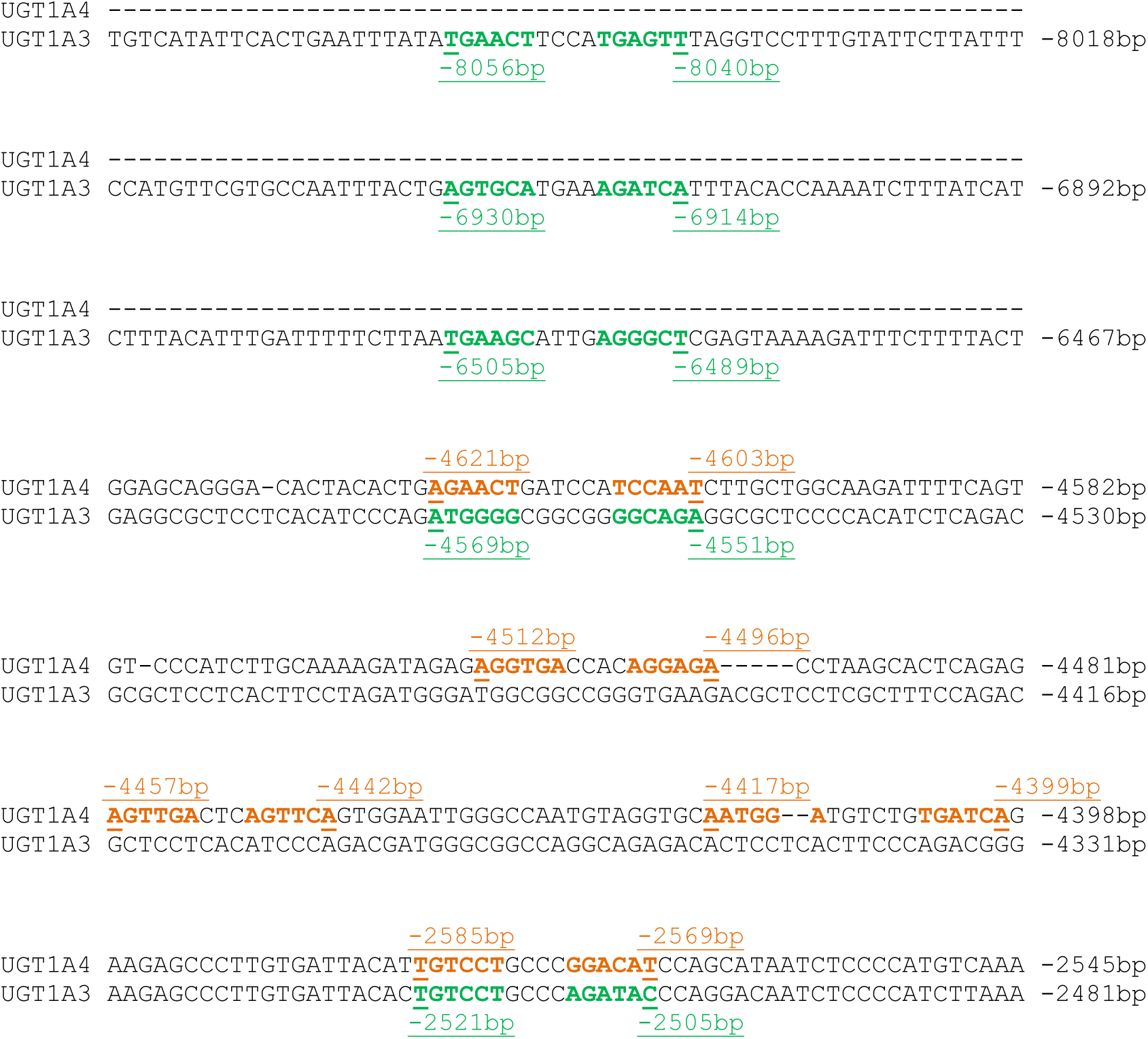
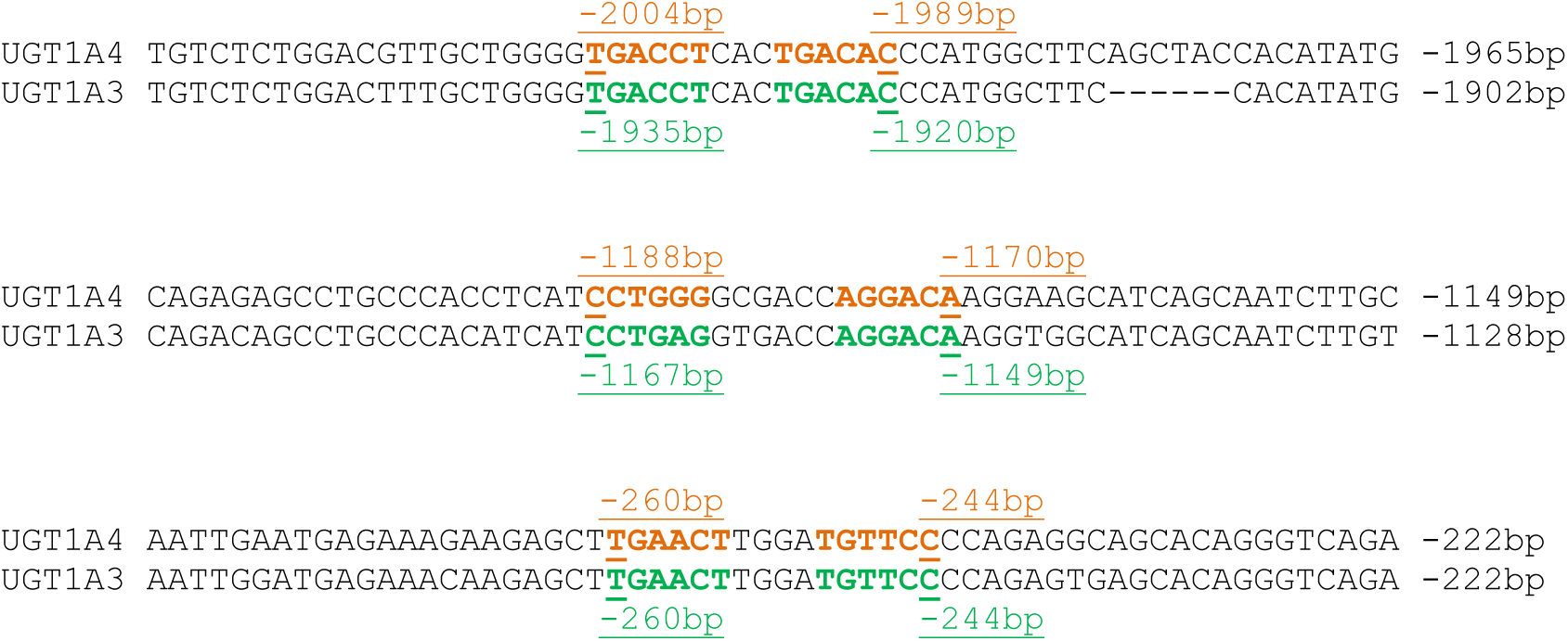

## Supplemental material 15. EMSAs performed with the DNA motifs from the UGT1A3 and 1A4 promoter genes identified as potential PXR response elements (PXRE)s.

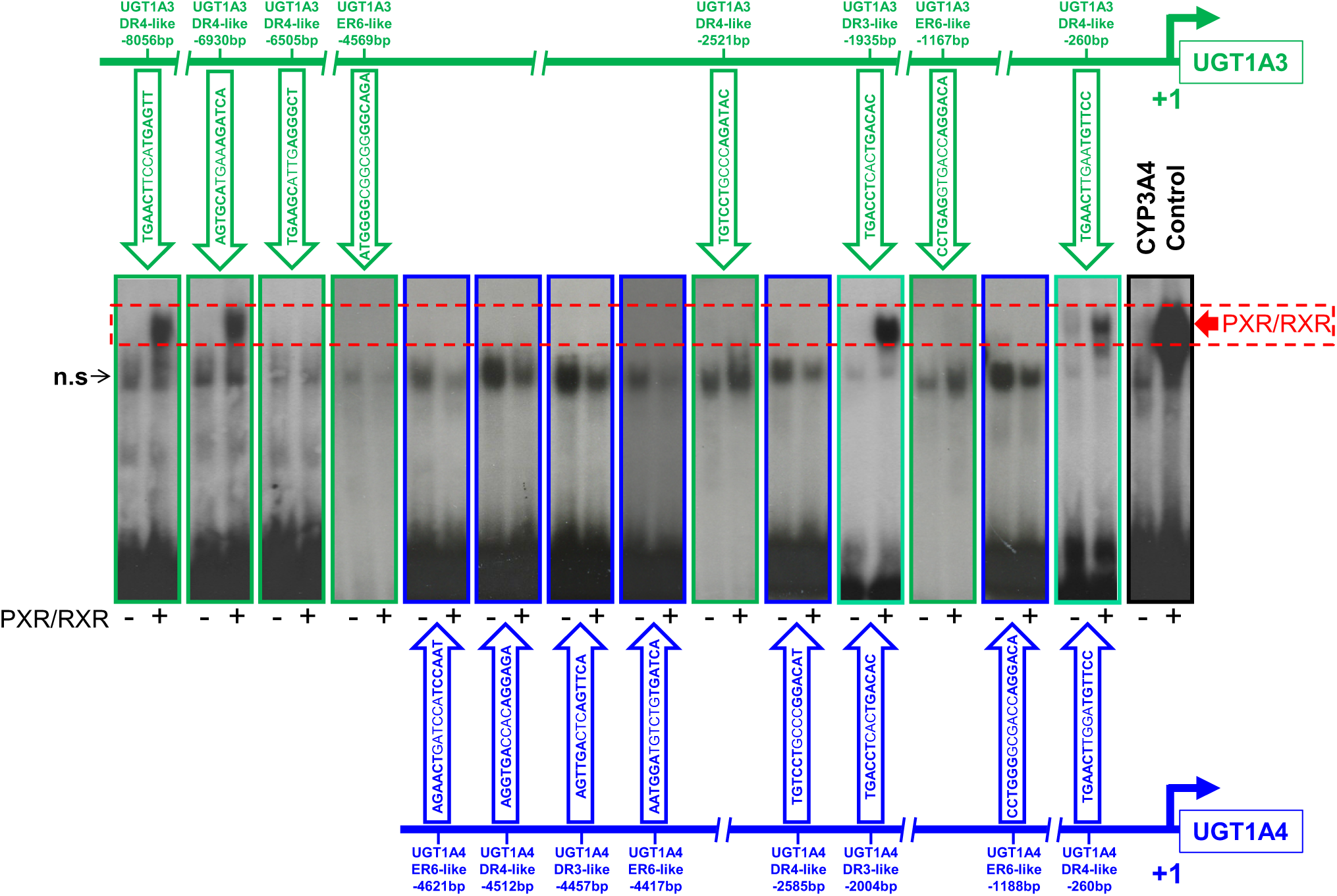

Electrophoretic Mobility Shift Assays (EMSA) were performed with the end-labeled DNA probes (100,000 cpm) listed in SM9 in the absence (-) or presence (+) of *in vitro* produced PXR and RXR proteins as indicated. Positions and sequences of these potential PXREs (SM14) are indicated. The previously reported(34) PXR-binding sequence from the human CYP3A4 gene has been used as a positive control.

## REFERENCES

1. McNally BB, Carey EJ. Cholestatic liver diseases: modern therapeutics. Expert Rev Gastroenterol Hepatol 2025;19:365–370.

2. Cancado GGL, Lleo A, Levy C, Trauner M, Hirschfield GM. Primary biliary cholangitis and the narrowing gap towards optimal disease control. Lancet Gastroenterol Hepatol 2025.

3. Manns MP, Bergquist A, Karlsen TH, Levy C, Muir AJ, Ponsioen C, Trauner M, et al. Primary sclerosing cholangitis. Nat Rev Dis Primers 2025;11:17.

4. Trottier J, Bialek A, Caron P, Straka RJ, Heathcote J, Milkiewicz P, Barbier O. Metabolomic profiling of 17 bile acids in serum from patients with primary biliary cirrhosis and primary sclerosing cholangitis: a pilot study. Dig Liver Dis 2012;44:303–310.

5. Trottier J, Bialek A, Caron P, Straka RJ, Milkiewicz P, Barbier O. Profiling circulating and urinary bile acids in patients with biliary obstruction before and after biliary stenting. PLoS One 2011;6:e22094.

6. Wunsch E, Trottier J, Milkiewicz M, Raszeja-Wyszomirska J, Hirschfield GM, Barbier O, Milkiewicz P. Prospective evaluation of ursodeoxycholic acid withdrawal in patients with primary sclerosing cholangitis. Hepatology 2014;60:931–940.

7. Li T, Chiang JYL. Bile acids as metabolic regulators: an update. Curr Opin Gastroenterol 2023;39:249–255.

8. Chiang JYL. New drug therapies for metabolic dysfunction-associated steatohepatitis. Liver Res 2025;9:94–103.

9. Tang Y, Fu A, Wang L, Ge Q. Microbiota-dependent metabolites - New engine for T cell warriors. Gut Microbes 2025;17:2523815.

10. Monte MJ, Marin JJ, Antelo A, Vazquez-Tato J. Bile acids: chemistry, physiology, and pathophysiology. World J Gastroenterol 2009;15:804–816.

11. Perreault M, Bialek A, Trottier J, Verreault M, Caron P, Milkiewicz P, Barbier O. Role of Glucuronidation for Hepatic Detoxification and Urinary Elimination of Toxic Bile Acids during Biliary Obstruction. PLoS One 2013;8:e80994.

12. Sharma R, Majer F, Peta VK, Wang J, Keaveney R, Kelleher D, Long A, et al. Bile acid toxicity structure-activity relationships: correlations between cell viability and lipophilicity in a panel of new and known bile acids using an oesophageal cell line (HET-1A). Bioorg Med Chem 2010;18:6886–6895.

13. Chatterjee S, Bijsmans IT, van Mil SW, Augustijns P, Annaert P. Toxicity and intracellular accumulation of bile acids in sandwich-cultured rat hepatocytes: role of glycine conjugates. Toxicol In Vitro 2014;28:218–230.

14. Paumgartner G, Pusl T. Medical treatment of cholestatic liver disease. Clin Liver Dis 2008;12:53–80.

15. Halilbasic E, Baghdasaryan A, Trauner M. Nuclear receptors as drug targets in cholestatic liver diseases. Clin Liver Dis 2013;17:161–189.

16. Trottier J, Milkiewicz P, Kaeding J, Verreault M, Barbier O. Coordinate regulation of hepatic bile acid oxidation and conjugation by nuclear receptors. Mol Pharm 2006;3:212–222.

17. Zhang Q, Lu L, Wang J, Lu M, Liu D, Zhou C, Liu Z. Metabolomic profiling reveals the step-wise alteration of bile acid metabolism in patients with diabetic kidney disease. Nutr Diabetes 2024;14:85.

18. Yang X, Wu R, Qi D, Fu L, Song T, Wang Y, Bian Y, et al. Profile of Bile Acid Metabolomics in the Follicular Fluid of PCOS Patients. Metabolites 2021;11.

19. Li W, Gong X, Niu X, Zhou Y, Ren L, Man Z, Tu P, et al. Quantitative comparison of bile acid glucuronides sub-metabolome between intrahepatic cholestasis and healthy pregnant women. Anal Bioanal Chem 2025;417:2823–2835.

20. Gallucci GM, Hayes CM, Boyer JL, Barbier O, Assis DN, Ghonem NS. PPAR-Mediated Bile Acid Glucuronidation: Therapeutic Targets for the Treatment of Cholestatic Liver Diseases. Cells 2024;13.

21. Trottier J, Perreault M, Rudkowska I, Levy C, Dallaire-Theroux A, Verreault M, Caron P, et al. Profiling serum bile acid glucuronides in humans: gender divergences, genetic determinants, and response to fenofibrate. Clin Pharmacol Ther 2013;94:533–543.

22. Kastrinou Lampou V, Poller B, Huth F, Fischer A, Kullak-Ublick GA, Arand M, Schadt HS, et al. Novel insights into bile acid detoxification via CYP, UGT and SULT enzymes. Toxicol In Vitro 2023;87:105533.

23. Trottier J, Verreault M, Grepper S, Monte D, Belanger J, Kaeding J, Caron P, et al. Human UDP-glucuronosyltransferase (UGT)1A3 enzyme conjugates chenodeoxycholic acid in the liver. Hepatology 2006;44:1158–1170.

24. Barbier O, Duran-Sandoval D, Pineda-Torra I, Kosykh V, Fruchart JC, Staels B. Peroxisome proliferator-activated receptor alpha induces hepatic expression of the human bile acid glucuronidating UDP-glucuronosyltransferase 2B4 enzyme. J Biol Chem 2003;278:32852–32860.

25. Gallucci GM, Trottier J, Hemme C, Assis DN, Boyer JL, Barbier O, Ghonem NS. Adjunct Fenofibrate Up-regulates Bile Acid Glucuronidation and Improves Treatment Response For Patients With Cholestasis. Hepatol Commun 2021;5:2035–2051.

26. Barbier O, Torra IP, Sirvent A, Claudel T, Blanquart C, Duran-Sandoval D, Kuipers F, et al. FXR induces the UGT2B4 enzyme in hepatocytes: a potential mechanism of negative feedback control of FXR activity. Gastroenterology 2003;124:1926–1940.

27. Erichsen TJ, Aehlen A, Ehmer U, Kalthoff S, Manns MP, Strassburg CP. Regulation of the human bile acid UDP-glucuronosyltransferase 1A3 by the farnesoid X receptor and bile acids. J Hepatol 2010;52:570–578.

28. Zhou X, Cao L, Jiang C, Xie Y, Cheng X, Krausz KW, Qi Y, et al. PPARalpha-UGT axis activation represses intestinal FXR-FGF15 feedback signalling and exacerbates experimental colitis. Nat Commun 2014;5:4573.

29. Trottier J, Caron P, Straka RJ, Barbier O. Profile of serum bile acids in noncholestatic volunteers: gender-related differences in response to fenofibrate. Clin Pharmacol Ther 2011;90:279–286.

30. Lai CQ, Arnett DK, Corella D, Straka RJ, Tsai MY, Peacock JM, Adiconis X, et al. Fenofibrate effect on triglyceride and postprandial response of apolipoprotein A5 variants: the GOLDN study. Arterioscler Thromb Vasc Biol 2007;27:1417–1425.

31. Wojczynski MK, Gao G, Borecki I, Hopkins PN, Parnell L, Lai CQ, Ordovas JM, et al. ApoB genetic variants modify the response to fenofibrate: a GOLDN study. J Lipid Res 2010;51:3316–3323.

32. Aron JH, Bowlus CL. The immunobiology of primary sclerosing cholangitis. Semin Immunopathol 2009;31:383–397.

33. Sumida A, Kinoshita K, Fukuda T, Matsuda H, Yamamoto I, Inaba T, Azuma J. Relationship between mRNA levels quantified by reverse transcription-competitive PCR and metabolic activity of CYP3A4 and CYP2E1 in human liver. Biochem Biophys Res Commun 1999;262:499–503.

34. Liu FJ, Song X, Yang D, Deng R, Yan B. The far and distal enhancers in the CYP3A4 gene co-ordinate the proximal promoter in responding similarly to the pregnane X receptor but differentially to hepatocyte nuclear factor-4alpha. Biochem J 2008;409:243–250.

35. Senekeo-Effenberger K, Chen S, Brace-Sinnokrak E, Bonzo JA, Yueh MF, Argikar U, Kaeding J, et al. Expression of the human UGT1 locus in transgenic mice by 4-chloro-6-(2,3-xylidino)-2-pyrimidinylthioacetic acid (WY-14643) and implications on drug metabolism through peroxisome proliferator-activated receptor alpha activation. Drug Metab Dispos 2007;35:419–427.

36. Verreault M, Senekeo-Effenberger K, Trottier J, Bonzo JA, Belanger J, Kaeding J, Staels B, et al. The liver X-receptor alpha controls hepatic expression of the human bile acid-glucuronidating UGT1A3 enzyme in human cells and transgenic mice. Hepatology 2006;44:368–378.

37. Maloney PR, Parks DJ, Haffner CD, Fivush AM, Chandra G, Plunket KD, Creech KL, et al. Identification of a chemical tool for the orphan nuclear receptor FXR. J Med Chem 2000;43:2971–2974.

38. Lehmann JM, McKee DD, Watson MA, Willson TM, Moore JT, Kliewer SA. The human orphan nuclear receptor PXR is activated by compounds that regulate CYP3A4 gene expression and cause drug interactions. J Clin Invest 1998;102:1016–1023.

39. Quandt K, Frech K, Karas H, Wingender E, Werner T. MatInd and MatInspector: new fast and versatile tools for detection of consensus matches in nucleotide sequence data. Nucleic Acids Res 1995;23:4878–4884.

40. Vyhlidal CA, Rogan PK, Leeder JS. Development and refinement of pregnane X receptor (PXR) DNA binding site model using information theory: insights into PXR-mediated gene regulation. J Biol Chem 2004;279:46779–46786.

41. Allen K, Jaeschke H, Copple BL. Bile acids induce inflammatory genes in hepatocytes: a novel mechanism of inflammation during obstructive cholestasis. Am J Pathol 2011;178:175–186.

42. Qin P, Borges-Marcucci LA, Evans MJ, Harnish DC. Bile acid signaling through FXR induces intracellular adhesion molecule-1 expression in mouse liver and human hepatocytes. Am J Physiol Gastrointest Liver Physiol 2005;289:G267–273.

43. Lindblad L, Lundholm K, Schersten T. Bile acid concentrations in systemic and portal serum in presumably normal man and in cholestatic and cirrhotic conditions. Scand J Gastroenterol 1977;12:395–400.

44. Setchell KD, Rodrigues CM, Clerici C, Solinas A, Morelli A, Gartung C, Boyer J. Bile acid concentrations in human and rat liver tissue and in hepatocyte nuclei. Gastroenterology 1997;112:226–235.

45. Song P, Zhang Y, Klaassen CD. Dose-response of five bile acids on serum and liver bile Acid concentrations and hepatotoxicty in mice. Toxicol Sci 2011;123:359–367.

46. Parks DJ, Blanchard SG, Bledsoe RK, Chandra G, Consler TG, Kliewer SA, Stimmel JB, et al. Bile acids: natural ligands for an orphan nuclear receptor. Science 1999;284:1365–1368.

47. Xie W, Radominska-Pandya A, Shi Y, Simon CM, Nelson MC, Ong ES, Waxman DJ, et al. An essential role for nuclear receptors SXR/PXR in detoxification of cholestatic bile acids. Proc Natl Acad Sci U S A 2001;98:3375–3380.

48. Zhang Y, Hong JY, Rockwell CE, Copple BL, Jaeschke H, Klaassen CD. Effect of bile duct ligation on bile acid composition in mouse serum and liver. Liver Int 2012;32:58–69.

49. Zollner G, Wagner M, Fickert P, Silbert D, Gumhold J, Zatloukal K, Denk H, et al. Expression of bile acid synthesis and detoxification enzymes and the alternative bile acid efflux pump MRP4 in patients with primary biliary cirrhosis. Liver Int 2007;27:920–929.

50. Boyer JL, Trauner M, Mennone A, Soroka CJ, Cai SY, Moustafa T, Zollner G, et al. Upregulation of a basolateral FXR-dependent bile acid efflux transporter OSTalpha-OSTbeta in cholestasis in humans and rodents. Am J Physiol Gastrointest Liver Physiol 2006;290:G1124–1130.

51. Zollner G, Fickert P, Silbert D, Fuchsbichler A, Marschall HU, Zatloukal K, Denk H, et al. Adaptive changes in hepatobiliary transporter expression in primary biliary cirrhosis. J Hepatol 2003;38:717–727.

52. Ros JE, Libbrecht L, Geuken M, Jansen PL, Roskams TA. High expression of MDR1, MRP1, and MRP3 in the hepatic progenitor cell compartment and hepatocytes in severe human liver disease. J Pathol 2003;200:553–560.

53. Oswald M, Kullak-Ublick GA, Paumgartner G, Beuers U. Expression of hepatic transporters OATP-C and MRP2 in primary sclerosing cholangitis. Liver 2001;21:247–253.

54. Bhushan S, Sohal A, Kowdley KV. Primary Biliary Cholangitis and Primary Sclerosing Cholangitis Therapy Landscape. Am J Gastroenterol 2025;120:151–158.

55. Eaton JE, Talwalkar JA. Primary Sclerosing Cholangitis: Current and Future Management Strategies. Curr Hepat Rep 2013;12:28–36.

56. Milkiewicz P, Wunsch E. Primary sclerosing cholangitis. Recent Results Cancer Res 2011;185:117–133.

57. Kowdley KV, Bowlus CL, Levy C, Akarca US, Alvares-da-Silva MR, Andreone P, Arrese M, et al. Efficacy and Safety of Elafibranor in Primary Biliary Cholangitis. N Engl J Med 2024;390:795–805.

58. Hirschfield GM, Bowlus CL, Mayo MJ, Kremer AE, Vierling JM, Kowdley KV, Levy C, et al. A Phase 3 Trial of Seladelpar in Primary Biliary Cholangitis. N Engl J Med 2024;390:783–794.

59. Caron P, Trottier J, Verreault M, Belanger J, Kaeding J, Barbier O. Enzymatic production of bile Acid glucuronides used as analytical standards for liquid chromatography-mass spectrometry analyses. Mol Pharm 2006;3:293–302.

60. Trottier J, El Husseini D, Perreault M, Paquet S, Caron P, Bourassa S, Verreault M, et al. The human UGT1A3 enzyme conjugates norursodeoxycholic acid into a C23-ester glucuronide in the liver. J Biol Chem 2010;285:1113–1121.

